# Quantifying stability via count splitting to guide model selection in RNA velocity analyses

**DOI:** 10.1101/2024.11.23.625009

**Authors:** Yuhong Li, Zeyu Jerry Wei, Yen-Chi Chen, Kevin Z. Lin

## Abstract

**Motivation:** RNA velocity is a computational framework that predicts future cell states from single-cell RNA sequencing data, offering valuable insights into dynamic biological processes. However, there is a lack of general methods to quantify the uncertainty and stability of these predictions from various RNA velocity methods.

**Results:** We present a novel framework for evaluating RNA velocity stability using negative binomial count splitting to generate independent data replicates, a metric we call *replicate coherence*. Testing five RNA velocity methods across datasets for mouse erythroid, pancreatic, and human brain development, we identify significant performance differences and inconsistencies, such as reversed velocity flows. Our framework remains robust even when intermediary cell states are missing. Furthermore, we introduce a signal-to-random coherence metric to guide model selection. We demonstrate that selecting fits with high replicate coherence uncovers more biologically informative gene pathways. This broadly applicable approach provides a rigorous tool for assessing and comparing RNA velocity methods across diverse biological contexts.

**Availability and implementation:** The code and analyses are available at https://github.com/linnykos/veloUncertainty.

## 1. Introduction

RNA velocity is a widely used computational tool in single-cell RNA sequencing (scRNA-seq) analyses, providing new insights into temporal transcriptomic dynamics (La Manno et al., 2018). By modeling the rate of change of the number of spliced and unspliced mRNA transcripts via an ordinary differential equation (ODE), these methods estimate high-dimensional velocity vectors for individual cells, capturing the direction of their developmental trajectories. When projected onto a 2D embedding, these velocity vectors enable the visualization of the entire cell differentiation process. The RNA velocity framework has been instrumental in making critical discoveries about the precise order of intermediary cell states in systems such as embryogenesis (Qiu et al., 2022) and lung regeneration (Lange et al., 2022). This importance has fueled the development of numerous follow-up RNA velocity methods, each making distinct assumptions and employing different statistical models. Although these RNA velocity methods exist to predict cell development, each can yield different results, making it difficult to assess their reliability and the uncertainty inherent in their predictions (Marot-Lassauzaie et al., 2022; Gorin et al., 2022; Bergen et al., 2021; Zheng et al., 2023). While multiple benchmark studies have compared different RNA velocity methods (Ancheta et al., 2024; Luo et al., 2025), the overarching conclusion is that no single RNA velocity method is universally superior. Hence, there is an urgent need for approaches to guide the selection of the most appropriate RNA velocity method for a given dataset.

The most commonly used metric for assessing an RNA velocity method’s performance is to measure the coherence of velocity vector estimates within neighborhoods and provide a confidence score, as demonstrated by Bergen et al. (2020). This metric works generically for any RNA velocity method. By computing the mean correlation between each cell’s velocity vector and its neighbors, the score measures the smoothness of the velocity vector field. While this metric quantifies the local agreement among the velocity vector estimates, it does not adequately evaluate the uncertainty of an RNA velocity method. This is because neighboring cells with similar transcriptomic profiles tend to exhibit similar velocity vectors, resulting in naturally high confidence scores. As such, even though this metric indicates local smoothness and coherence, it does not reflect the uncertainty inherent in the method’s output (Ancheta et al., 2024). Additionally, although other metrics quantify RNA velocity uncertainty, they are often specific to a particular RNA velocity method. For example, VeloVI (Gayoso et al., 2024) uses variational inference and introduces intrinsic and extrinsic uncertainty computations based on sampling from the posterior of their fitted Bayesian model.

These limitations motivate us to devise a framework for quantifying uncertainty in RNA velocity methods that is not confounded by cell local similarity and is independent of the specific method used. Our approach leverages negative binomial count splitting (Neufeld et al., 2023), which decomposes any count matrix into multiple statistically independent partitions. This procedure can be interpreted as creating independent replicates of each cell, enabling the comparison of a cell to its replicate, as if the cell were observed multiple times along the biological trajectory. This idea has been recently used to assess the clustering stability of scRNA-seq data (Neufeld et al., 2024; Sant et al., 2025). Count splitting aims to create multiple independent *n* × *p* matrices (typically two) that entry-wise sum to the original matrix. The resulting splits can be interpreted as synthetic “technical replicates,” where we sequence the exact same set of cells multiple times. In reality, it is only possible to sequence the same cell once due to the destructive nature of single-cell sequencing. Nonetheless, this statistical framework allows us to compare the behavior of the same cells and quantify the stability of different RNA velocity methods across synthetic, independent replicates. We will refer to velocity confidence defined in Bergen et al. (2020) as *local coherence* and our uncertainty measure as *replicate coherence* in the following text.

## 2. Methods

### 2.1 Replicate coherence via count splitting

We present a framework for evaluating *replicate coherence*, which leverages negative binomial count splitting (Neufeld et al., 2023) to create two independent splits of scRNA-seq data and assess the stability of RNA velocity estimates, providing a way to compare the performance of different velocity methods (Figure 1a).

**Fig. 1:**
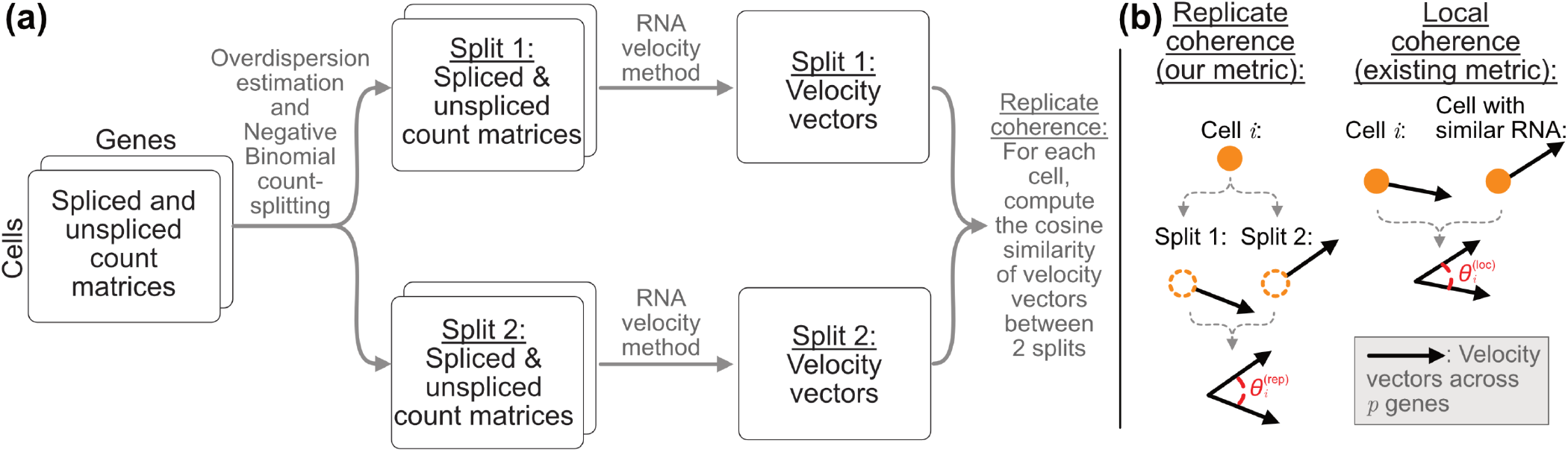
Diagram illustrating replicate coherence. **(a)** Starting with the initial spliced/unspliced single-cell RNA-sequencing count matrices, we estimate the overdispersion parameter and perform negative binomial count splitting on both matrices to obtain two sets of spliced and unspliced datasets. We then run an RNA velocity method on each split separately, obtaining two sets of cell-specific velocity vectors. The replicate coherence for each cell is computed as the cosine similarity between its two RNA velocity vectors, one from each of the two splits. **(b)** Illustration showcasing the difference between replicate coherence 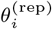 and local coherence 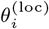 for cell *i*.

We denote the single-cell RNA-sequencing count matrix with *n* cells and *p* genes as 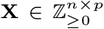, which can denote either the spliced or unspliced count matrix. Assume that the expression count of the *j*-th gene in cell *i* follows a negative binomial distribution, **X**_*ij*_ ∼ NB(*µ*_*ij*_, *γ*_*j*_), with mean 𝔼 [**X**_*ij*_] = *µ*_*ij*_ and variance 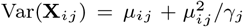, where each *γ*_*j*_ *>* 0 denotes a gene-specific overdispersion parameter (Sarkar and Stephens, 2021). A finite non-negative *γ*_*j*_ indicates overdispersion, where the variance exceeds the mean, and an infinite *γ*_*j*_ indicates the distribution approaches a Poisson distribution with mean *µ*_*ij*_ . For each gene, we fit a Gamma-Poisson Generalized Linear Model (GLM) without covariates to estimate its overdispersion parameter for the count splitting procedure. In our analysis, each dataset consists of a spliced and unspliced data matrix, and we estimate the overdispersion parameters for the two matrices separately.

We then apply negative binomial count splitting to generate two splits **X**^(1)^, **X**^(2)^, such that **X**^(1)^ + **X**^(2)^ = **X**. For each entry,

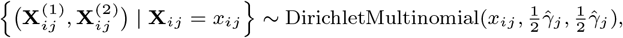

where 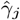 _*j*_ is the estimated overdispersion parameter for gene *j*. Here, DirichletMultinomial(*n, p*_1_, *p*_2_) draws a probability vector *p* specified by Dirichlet(*p*_1_, *p*_2_), and then samples the *n* counts via a multinomial distribution to split 1 or 2 with probabilities specified by *p*. Neufeld et al. (2023) proved the theoretical properties of count splitting that: 1) the splits **X**^(1)^, **X**^(2)^ are independent, and 2) both 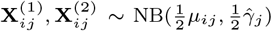 for each cell *i* and gene *j*. The first condition ensures that **X**^(1)^ and **X**^(2)^ can be treated as “technical replicates,” with estimation performance in one split independent of the other. The second condition ensures that both splits constitute valid scRNA-seq datasets and are suitable for standard single-cell analysis workflows.

We note that the independence guarantee relies on using the true overdispersion parameters, and plug-in estimates can induce dependence between splits. Empirically, however, the splits are nearly uncorrelated based on the estimated overdispersion, and this slight dependence does not materially affect count splitting’s performance (see Supplementary Figures S1, S3).

After count splitting, we get two independent count matrices that sum to the original counts, each preserving a negative binomial distribution. We run the same RNA velocity method on each split separately. An overview of how each method estimates the RNA velocity via ordinary differential equation modeling is provided in Supplementary Section S1. For methods requiring separate spliced and unspliced matrices, we apply count splitting to each matrix, yielding two spliced count matrices and two unspliced count matrices. Applying an RNA velocity method to each split produces two sets of cell-specific velocity matrices **V**^(1)^, **V**^(2)^ ∈ ℝ^*n*×*p*^ for **X**^(1)^, **X**^(2)^ respectively, where the *i*-th row 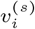 represents the velocity vector for cell *i* across all *p* genes based in split *s* ∈ {1, 2}. To assess the coherence between these two results, we compare the cell-specific velocity vectors from the two splits and measure the alignment for cell *i* using cosine similarity,

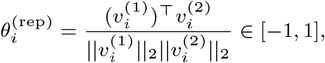

where ∥· ∥_2_ denotes the Euclidean norm. We call this quantity the *replicate coherence* of cell *i*. Values close to +1 indicate that the two velocity vectors point in similar directions, suggesting a stable velocity estimate, while values near −1 reflect opposite directions and divergent results.

A popular alternative for measuring the “confidence” of an RNA velocity fit is *local coherence*, as initially proposed by scVelo (Bergen et al., 2020). This metric compares each cell’s velocity estimate to those of its neighbors. Specifically, let *v*_*i*_ denote the velocity vector of cell *i*, ne(*i*) denote the set of neighboring cells of cell *i*, and *p* denote the number of genes in each velocity vector. Further, let 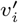 denote the centered velocity vector for cell *i*, where each element 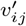 is defined as 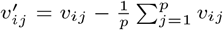. The local coherence for cell *i* is then computed as

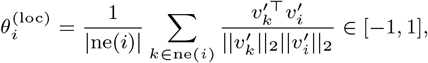

which is essentially the mean cosine similarity between a cell’s centered velocity vector and those of its neighbors, capturing the local smoothness. Both replicate and local coherence provide useful measures of RNA velocity fit, but local coherence can be artificially inflated: because it is computed after smoothing gene expression across cell neighborhoods, neighboring velocity vectors are similar by construction. Replicate coherence avoids this issue by comparing velocity estimates from independent data splits, providing a more methodologically sound measure of method stability. The distinction between the two metrics is illustrated in Figure 1b.

We include five different RNA velocity methods in our work: scVelo (Bergen et al., 2020), UniTVelo (Gao et al., 2022), scTour (Li, 2023), the standard VeloVI with the preprocessing step, and VeloVI without the preprocessing step (Gayoso et al., 2024). Briefly, scVelo and VeloVI both model spliced and unspliced mRNA counts, with scVelo estimating gene-specific kinetics and VeloVI providing uncertainty quantification. UniTVelo uses a spliced-only approach, utilizing a unified latent time across genes, whereas scTour models total RNA counts and projects velocities onto a low-dimensional space. We evaluate the five RNA velocity methods on three scRNA-seq datasets: mouse erythroids during gastrulation (Pijuan-Sala et al., 2019), mouse pancreatic endocrinogenesis development (Bastidas-Ponce et al., 2019), and human brain development (Trevino et al., 2021). Details of the datasets are given in Supplementary Section S2. In Supplementary Section S3, we document how we compute the replicate coherence for each method since some differ in their inputs or outputs. Importantly, our framework involves comparing high-dimensional velocity vectors, but methods like scTour output the RNA velocity in a low-dimensional latent embedding space. In Supplementary Section S3, we document how we handle scenarios like this and demonstrate its validity in Supplementary Figure S16.

### 2.2. Robustness diagnostics for intrinsic randomness and missing cell types

With the framework and replicate coherence in place, we can evaluate the robustness of an RNA velocity method. Firstly, the inherent randomness of count splitting can introduce variability across different random seeds. To mitigate this, we repeat the procedure using multiple seeds and demonstrate the robustness of our results. In practice, if resources permit, the computation procedure can be repeated five or more times to assess a method’s stability.

Secondly, in real-world settings, intermediate cell states along the developmental trajectory may be underrepresented due to factors such as short lifespans, low abundance, or technological limitations. Prior work on RNA velocity has raised concerns about this issue (Atta et al., 2022; Li, 2023; Zheng et al., 2023). Inspired by Li (2023), we investigate how replicate coherence behaves when intermediate cell states are omitted, with the expectation that a reliable quantification of cellular velocity should remain robust even in the absence of such cell states. We first apply RNA velocity to the full dataset, then remove an intermediate cell type and reanalyze, comparing the method’s replicate coherence across the two settings.

### 2.3. Signal-to-random coherence score to aid in selecting an RNA velocity fit

For a given dataset, we also propose a metric to guide RNA velocity method selection which captures how well a method distinguishes datasets with strong biological signals from those with weak or irrelevant signals. We expect a reliable method to behave differently in these two settings. To assess this, we construct two gene subsets: *informative genes* with the most relevance to the biology of interest and *non-informative genes* with little relevance. We consider these two distinct gene sets since prior work has shown that analyzing too many genes across diverse pathways could obscure the biological process of interest (Lotfollahi et al., 2023; Ranek et al., 2024). We then compare replicate coherence from an RNA velocity method applied to each gene set. If replicate coherence is similar across both sets, it suggests that the inferred trajectories, regardless of how plausible they appear, may have arisen by chance rather than by biological signal and should be interpreted with caution.

As outlined above, the first step is to select informative and non-informative gene sets. Informative genes can be defined based on marker genes of the various cell types, a multi-omic analysis tracking how different modalities change during development, or biological knowledge from the literature. To construct a non-informative control set, we take inspiration from ChromVAR (Schep et al., 2017) and match genes to the informative set based on (1) spliced expression sparsity, (2) unspliced expression sparsity, and (3) gene-specific *R*^2^ values from scVelo which reflects the expression variability explained by the steady-state model. Each of these three features is divided into four bins, yielding 4^3^ = 64 groups, and each gene is assigned to a bin. Euclidean distances between bins are then computed. For each informative gene, a non-informative gene is randomly selected from the same bin that is not labeled informative and not already matched. If none is available in the same bin, we select from the nearest bin based on Euclidean distance. This ensures matched non-informative genes resemble informative genes in sparsity and *R*^2^, but differ in biological relevance. In practice, equal-sized gene sets often yield too few genes passing velocity preprocessing, sometimes causing errors in certain methods. To ensure fair comparison, we set the non-informative gene set to be twice as large as the informative gene set.

Next, we compute the *signal-to-random coherence score*, which measures replicate coherence on the informative gene set relative to the non-informative gene set. Since non-informative genes are randomly sampled, we compute the signal-to-random coherence score across *B* independent runs (i.e. *B* random seeds). Each random seed *b* ∈ {1, …, *B*} for an RNA velocity method requires: (1) an independent run of count splitting, (2) sampling non-informative genes, and (3) two runs of the RNA velocity method. Let 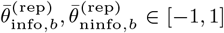 denote the mean replicate coherence across *n* cells for the informative and non-informative gene sets, respectively, at seed *b*. After *B* runs, the signal-to-random coherence score for an RNA velocity method is defined as

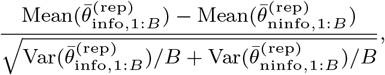

where 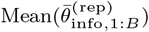 and 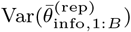 denote the mean and variance of replicate coherence across *B* seeds. This score is inspired by Welch’s t-test, but we use this score for method selection rather than for hypothesis testing.

After computing the signal-to-random coherence score for all methods, we recommend selecting the fit with the highest score. As shown in Section 3.3, this procedure identifies a biologically meaningful fit based on the relevant gene pathways. However, as it depends on the availability and choice of well-characterized informative and non-informative gene sets, which may not be feasible for every dataset, practitioners should assess whether this approach is appropriate for their context.

## 3. Results

### 3.1. Evaluation of replicate coherence in RNA velocity analyses

We use the mouse erythroid dataset (Pijuan-Sala et al., 2019) to demonstrate the validity of count splitting and the utility of our replicate coherence score. This dataset contains 9,815 cells across five cell types, where blood progenitor cells gradually differentiate into erythroid cells. Using this dataset, we demonstrate that our replicate coherence, introduced in Section 2.1, can help assess the robustness of results from an RNA velocity analysis. Our premise is that if we apply count splitting and apply the same RNA velocity workflow on each split, a misalignment between the two resulting RNA velocity fields indicates potential computational instability in the method.

We first validate the theoretical result of the independence among genes between the two splits. Among genes with non-zero counts in both splits, most exhibit correlations close to 0, supporting the notion of treating the two splits as technical replicates (see Supplementary Figure S1, S3). We provide additional investigations in Supplementary Figure S5, where we visualize the impact of count splitting in more detail and ensure that the overdispersion estimates are reliable by assessing their relationship to the sparsity of the genes.

Next, by running all five RNA velocity methods on the dataset, we find that UniTVelo, scTour, and VeloVI, without the preprocessing step, can correctly identify the trajectory (Figure 2a). In contrast, scVelo and standard VeloVI produce reversed predictions, with velocity flow pointing from erythroid cells back to blood progenitor cells. Such a pattern is not directly reflected in the measure of local coherence, where all methods except the standard VeloVI show high values close to 1, indicating the overall smoothness of velocity vectors within cell neighborhoods (Figure 2b). However, regarding replicate coherence, scVelo and standard VeloVI exhibit median values as low as 0.5, suggesting relative instability across technical replicates and corresponding to less precise performance. In contrast, the replicate coherence values of the other methods remain relatively high. We reiterate that our replicate coherence was computed directly based on the velocity vectors 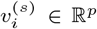 for each cell *i* between splits *s* ∈ {1, 2}. We emphasize this since work like Zheng et al. (2023) has emphasized the ambiguity of projecting RNA velocity vectors onto a UMAP embedding. The authors showed that the UMAP visualizations, such as those in Figure 2b, can sometimes be misleading. However, since our replicate coherence is computed based on the velocity vectors across all the genes, we provide a concrete and orthogonal metric to reinforce that the reversed predictions in scVelo and standard VeloVI are not artifacts of visualization.

**Fig. 2:**
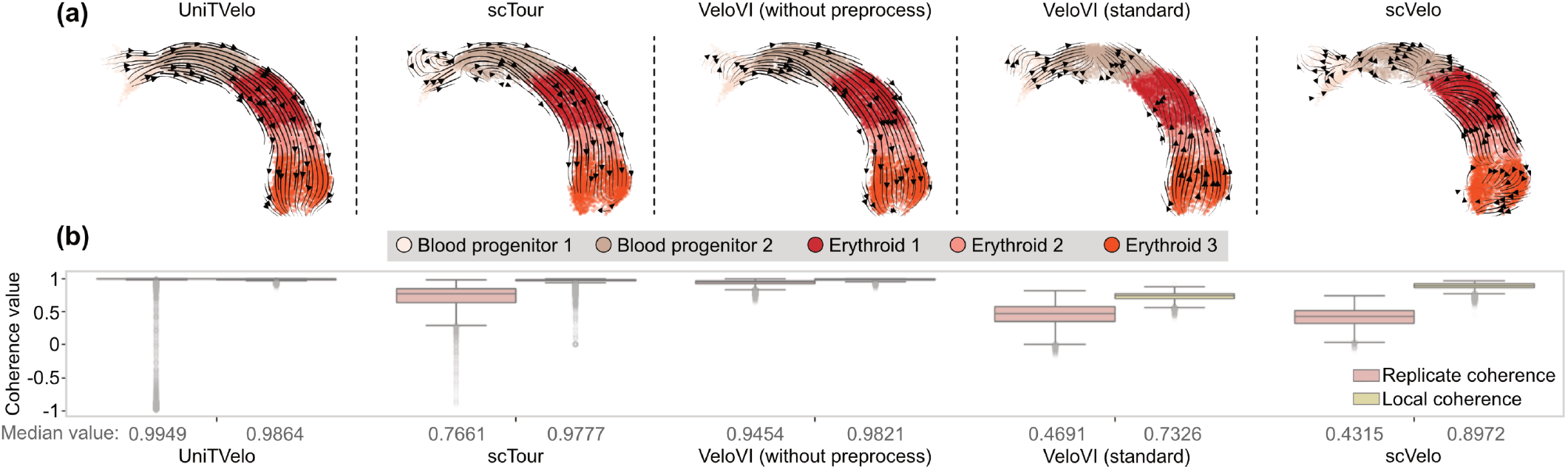
Evaluation of replicate coherence based on mouse gastrulation dataset (Pijuan-Sala et al., 2019). **(a)** The velocity estimates plotted on UMAPs for each of the five RNA velocity methods, with the embeddings directly inherited from the original dataset. **(b)** Boxplots of replicate or local coherence for each RNA velocity method, where we report the median replicate coherence across the *n* cells below the boxplots (red dashed line: coherence of 0).

Because count splitting yields two independent sets of spliced and unspliced count matrices, we can investigate the source of an RNA velocity method’s low replicate coherence in more detail. We demonstrate this conflicting behavior of scVelo based on the two splits in the erythroid dataset (Supplementary Figure S6). Note that we use the UMAPs computed from the split data rather than the UMAP from the entire dataset object, as shown in Figure 2b. In Split 1, unlike the behavior on the entire dataset, the predictions appear surprisingly reasonable. However, Split 2 shows a more disorganized pattern and deviates from both Split 1 and the true trajectory, resulting in low replicate coherence. The full details of our analysis can be found in Supplementary Section S3.

### 3.2. Robustness of replicate coherence with respect to intrinsic randomness and missing cell types

We use the mouse pancreas dataset (Bastidas-Ponce et al., 2019) to demonstrate how a method’s replicate coherence is preserved, even when cell types at intermediary developmental times are purposefully removed before the analysis, as we introduced in Section 2.2. This dataset contains 3696 cells across eight cell types, where, critically, pre-endocrine cells are a necessary intermediary cell state that cells must go through before becoming alpha, beta, delta, and epsilon cells (Figure 3a). Here, we visualize the RNA velocity estimate using scVelo as an example to mimic the original publication (Bergen et al., 2020).

**Fig. 3:**
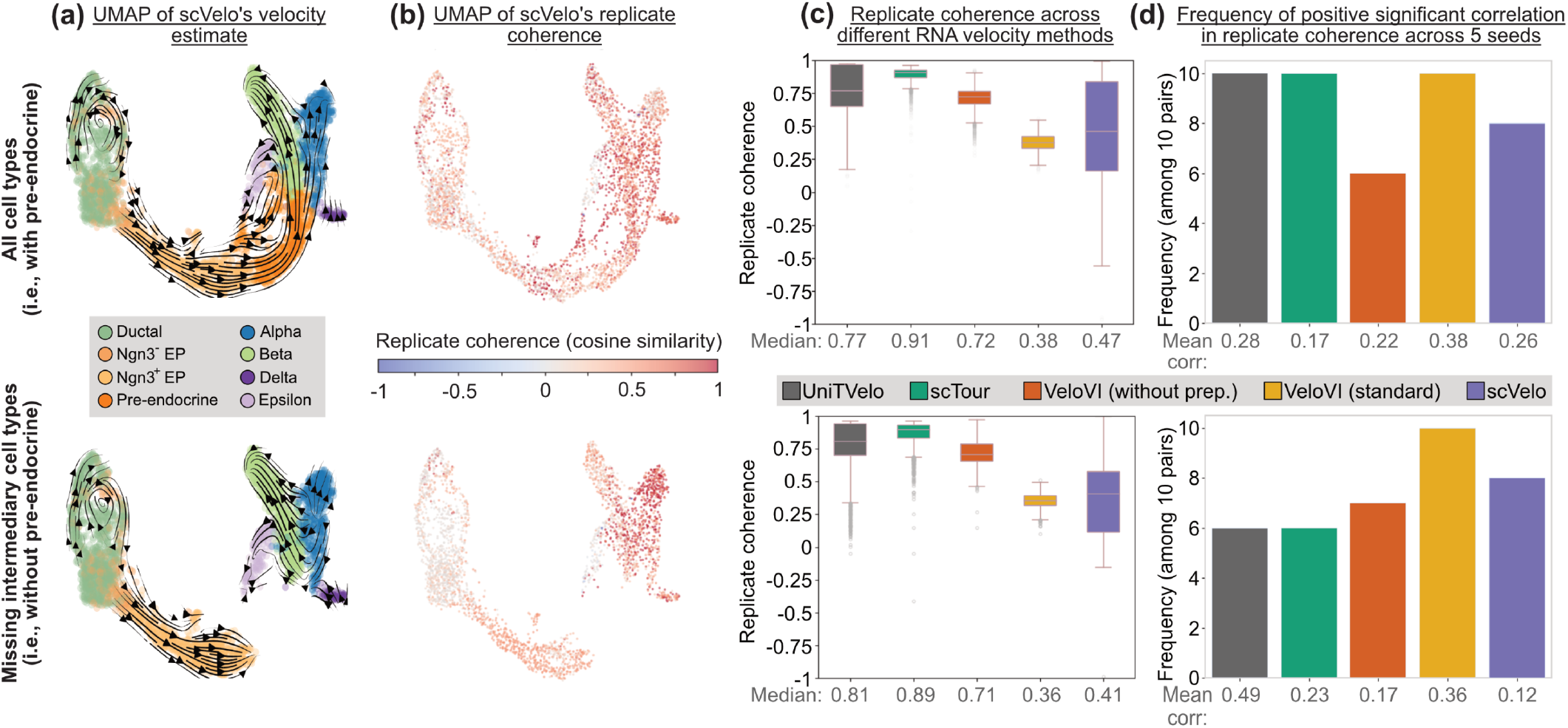
Evaluation of robustness of replicate coherence with respect to missing cell types and intrinsic randomness. **(a)** Velocity estimates of scVelo on UMAPs of pancreas dataset (Bastidas-Ponce et al., 2019) with and without pre-endocrine cells. **(b)** UMAPs coloring cells by replicate coherence values. **(c)** Boxplots showing replicate coherence across five RNA velocity methods across five RNA velocity methods on the pancreas data with and without pre-endocrine cells, with the median replicate coherence values indicated at the bottom. **(d)** Frequencies of positive correlations with significant p-values for replicate coherence correlations computed across five seeds for five RNA velocity methods, with the mean correlation reported at the bottom.

Visually, all five methods preserve similar predicted velocity flows on the UMAP embeddings in both datasets. For instance, Figure 3a,b display the RNA velocity estimates and replicate coherence from scVelo on the two datasets: the pancreas dataset with and without the pre-endocrine cells, respectively. The velocity estimates from the other four methods are in Supplementary Figure S9. Scanning across all the RNA velocity methods, we see that in Figure 3c that the replicate coherence is comparable in terms of the relative ordering of different methods across both datasets. Since the replicate coherence estimates are similar between the pancreas dataset with and without pre-endocrine cells, we demonstrate that this metric is robust to the presence of unobserved cell states. For comparison, we also report VeloVI’s intrinsic and extrinsic uncertainty quantification in Supplementary Figures S8 and S9. Additional plots visualizing the local and replicate coherence are in Supplementary Figures S11-S15.

Next, we examine whether the intrinsic randomness in generating the count matrix that count splitting introduces affects the reliability of our replicate coherence. To investigate this, we split the two pancreas datasets five times (i.e., five seeds), setting the random seed to a different value for each setting. We then apply each RNA velocity method to the five pairs of count matrices. We compute replicate coherence on each pair of splits for each seed within a method and test the significance of Pearson’s correlations for the replicate coherence values. In particular, since there are five different replicate coherence values for any specific RNA velocity method, we perform 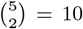 hypothesis tests to assess whether the correlation among the cell’s replicate coherence values between any two sets of results is significantly positive. Our results show that, for all methods, most correlations are positive and associated with significant p-values at a significance level of 0.05, using the Bonferroni correction (Figure 3d). Furthermore, all methods exhibited mean correlations well above 0. Additionally, in Supplementary Section 3, we show that the “uncentered” correlation is always significantly positive, demonstrating that the positivity of a method’s replicate coherence is not impacted by randomness. This indicates that despite the randomness in this framework, replicate coherence remains robust across different seeds for each method. The replicate coherence correlations across multiple seeds are plotted in pairs in Supplementary Figures S17-S21, Table S5. We performed only five seeds due to limitations in computational resources. See Supplementary Table S6 for details on the time required to run each method.

### 3.3. Signal-to-random coherence score to select a RNA velocity fit and downstream biological insights

Having established the replicate coherence that is shown to differentiate between the biologically correct versus reversed trajectories (Section 3.1) and robust (Section 3.2), we are now ready to demonstrate how the replicate coherence can aid in determining the most appropriate RNA velocity fit for any particular dataset. To illustrate that the signal-to-random coherence score is effective for this task, we showcase its application on a human brain development dataset (Trevino et al., 2021). This dataset contains more complex and differentiated trajectories compared to the previous datasets, comprising 38,262 cells that represent various stages of glial or neuronal differentiation. The analogous results, applying the signal-to-random coherence score to the mouse gastrulation and pancreas datasets, are shown in Supplementary Figures S22 and S23.

We apply the standard workflow to both informative and non-informative gene sets and compute the signal-to-random coherence score for five RNA velocity methods. The informative genes are selected as a set of 185 genes with predictive chromatin (GPC) determined by the original authors, where their expression is predictable from chromatin accessibility. Figure 4a presents the signal-to-random coherence score, where positive values indicate that the mean replicate coherence for informative genes exceeds that for non-informative genes. We observe that VeloVI without preprocessing achieves the largest positive score of 3.45. In contrast, scVelo yields the lowest negative score of −5, suggesting a notably poorer performance on informative genes relative to non-informative ones in terms of replicate coherence. To investigate the cells driving this large discrepancy, we compare the cell-specific velocity vectors pairwise, in terms of their angles (Figure 4b). The results show that, overall, Excitatory Neuron 4 cells exhibit larger angles between velocity vectors from the two methods, indicating substantial disagreement in their estimates for this cell type.

**Fig. 4:**
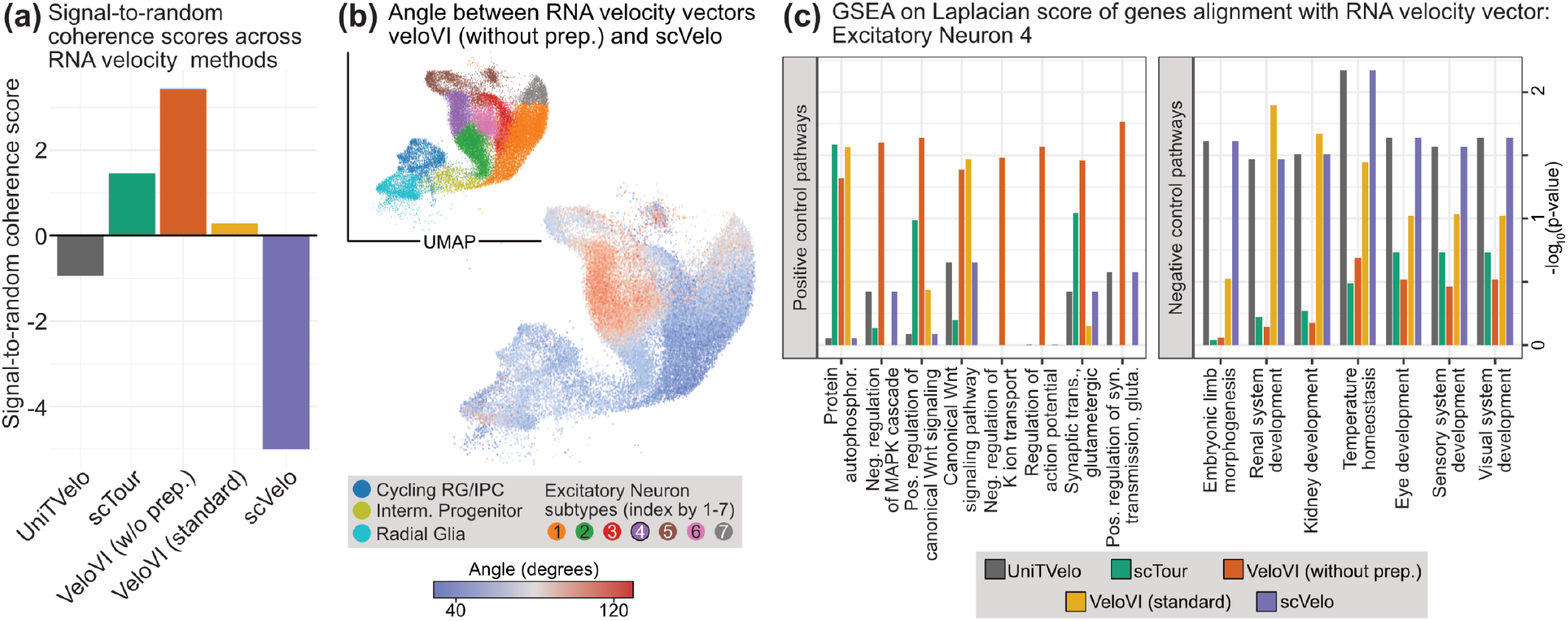
Signal-to-random coherence score selects biologically meaningful RNA velocity fits. **(a)** The signal-to-random coherence score across five RNA velocity methods analyzing the human brain development dataset (Trevino et al., 2021). A higher value (better) indicates an RNA velocity fit with a higher coherence score compared to a biologically uninformative analysis. **(b)** Angle difference in the RNA velocity fits between the fit from veloVI without preprocessing and scVelo as a UMAP, ranging from red (most opposite) to blue (most similar). The inset shows the cells annotated by cell types. **(c)** GSEA based on the direction-aware Laplacian transition matrix of “positive” and “negative” control pathways, focusing on the Excitatory Neuron 4 cells, where a high − log_10_ p-value for the pathway denotes a pathway more aligned with the estimated RNA velocity vectors.

Based on this analysis, we next investigate if we would derive drastically different biological insights regarding the Excitatory Neuron 4 cells based on these two RNA velocity estimates. We compute gene-specific scores that quantify how strongly each gene aligns with the transition matrix among these cells obtained by each RNA velocity method (Lange et al., 2022), and then apply a gene set enrichment analysis (GSEA) on these gene-specific scores. The exact details are documented in the Supplementary Section 3, where we explain how we construct a direction-aware Laplacian transition matrix (Chung, 2005).

To contextualize the GSEA results, we first identify so-called “positive” and “negative” control pathways based on the scientific literature exploring their biological association with the differentiation of excitatory neurons. For example, as seen in Figure 4c, some positive control pathways in our analysis involve the Wnt signaling pathway and the MAPK cascade which balance radial-glial proliferation and neuronal commitment (Da Silva et al., 2021). Additionally, pathways regarding synaptic transmission and action potential are known to be involved during excitatory neuron differentiation (Liang et al., 2021; Duque et al., 2013). In contrast, the negative control pathways involve ones primarily associated with other biological systems outside the brain. With this dichotomization of specific pathways, we observe that many positive control pathways are deemed more significant when using veloVI without preprocessing. In contrast, many negative control pathways related to the renal or visual sensory systems are deemed more significant when using scVelo, suggesting that these “random” pathways are correlated with the RNA velocity fit. While these negative control pathways are not definitively unrelated to the differentiation of excitatory neurons during embryogenesis, there is a lack of direct literature linking these two processes. Overall, our results provide evidence that meaningful biological pathways corroborate the high signal-to-random coherence score. We provide additional details on this analysis as well as analogous results on the erythroid and pancreas datasets (Supplementary Figures S24 and S25).

## 4. Discussion

In this paper, we present a framework to assess the stability of velocity predictions in RNA velocity methods. By creating two sets of hypothetical twin cells via count splitting, we compare the method’s results on two technical replicates of the same biological process, providing a measure of uncertainty. Furthermore, by examining how a method behaves on datasets with strong versus weak biological signals, we quantify its performance and use this score to select the method whose fit reflects biologically meaningful pathways. We apply our framework to five RNA velocity methods across three datasets, demonstrating its (i) effectiveness in distinguishing correct from incorrect method performance in mouse erythroid development, (ii) robustness to random seeds and unobserved time intervals using the pancreatic endocrinogenesis dataset, and (iii) usage in quantifying each method’s performance relative to the underlying biology. Notably, our framework offers a generic approach to measuring the stability of any RNA velocity method, unlike VeloVI’s intrinsic and extrinsic uncertainty.

In our work, replicate coherence is based on high-dimensional velocity vectors, which are typically the main output of an RNA velocity method. However, methods such as scVelo and UniTVelo internally select a subset of genes to model, so the exact set of genes used may differ between two splits. To address this, we compute cosine similarity using the union of genes across both splits, padding entries with 0 for genes present in one split but absent in the other. In this approach, all genes used in either split contribute to the computation, but we penalize the replicate coherence when genes are selected in only one split. Supplementary Table S2 summarizes the number of genes in the intersection and union of the gene sets in two splits for each method. Moreover, in our current analysis of replicate coherence, we primarily use boxplots and median values for qualitative comparisons. To support a quantitative evaluation, we also compare the standard replicate coherence to those obtained from shuffled cell pairs, shown in Supplementary Table S9. This procedure creates a null distribution to assess whether the observed cosine similarities exceed those expected under random cell pairing within a method.

One potential limitation of our current work is that we applied the framework to three well-characterized scRNA-seq datasets. However, real-world datasets often present greater complexity. We view our proposed replicate coherence as a core data-driven quality metric for selecting the most appropriate RNA velocity fit, but it is not necessarily sufficient by itself to select the most accurate fit. Because RNA velocity is an unobserved concept that depends on the assumed biological model (Gorin et al., 2022), it is difficult, without specialized technologies such as fluorescent ubiquitination-based cell-cycle indicator (FUCCI; Mahdessian et al. (2021)), to assess the accuracy of an RNA velocity method. Future work could test the framework on more diverse datasets that reflect real-world limitations, identify factors influencing performance, and refine the approach to handle these complexities effectively. Additionally, the signal-to-random coherence score provides a numerical summary but relies on the selection of gene sets with varying biological relevance. While informative, it is a descriptive metric rather than a formal statistical test. One direction for future work is to incorporate replicate coherence into a hypothesis testing framework, enabling statistical inference and providing more rigorous guidance for RNA velocity method selection. Finally, since the overdispersion parameters must be estimated in practice, exact independence between splits is not guaranteed. An alternative future research direction is to develop split-compare procedures that avoid plugging in additional parameters while still producing independence.

## Data availability

The mouse erythroid dataset (Pijuan-Sala et al., 2019) is from https://scvelo.readthedocs.io/en/stable/scvelo.datasets.gastrulation_erythroid.html, the mouse pancreatic endocrinogenesis dataset (Bastidas-Ponce et al., 2019) is from https://scvelo.readthedocs.io/en/stable/scvelo.datasets.pancreas.html, both from the scVelo (Bergen et al., 2020) codebase, and the human brain development dataset (Trevino et al., 2021) is from https://github.com/GreenleafLab/brainchromatin, where the links are provided in the links.txt file.

## Author contributions statement

YL, ZJW, YC, and KZL all contributed to the ideas behind this paper. YL and ZJW developed the code for the method. YL and KZL performed the analyses and wrote the paper.

## Acknowledgments and funding

We thank Lucy Gao, Dan Kessler, Anna Neufeld, and Daniela Witten for insightful discussions regarding count splitting. This work was supported by the Genentech Endowed Professorship and NIGMS of the NIH under award number 1R35GM162089.

## Supplementary data

The supplement provides more details, including visualizations and comparisons of our results and a simulation study. Additional plots of the variability across different random seeds, computation time, and the signal-to-random scores, pathways are also shown.

