## Supplementary Information for "Quantifying stability via count splitting to guide model selection in RNA velocity analyses"

### S1. Review of replicate coherence and RNA velocity methods

#### S1.1. Review of RNA velocity methods

***scVelo* (Bergen et al., 2020)**
ScVelo is a likelihood-based model of the dynamics of spliced and unspliced mRNA. The method estimates gene-specific kinetic parameters by solving an ODE system, including transcription, splicing, and degradation rates. There are two modes of velocity computation. The stochastic model accounts for the balance of spliced and unspliced mRNA counts as well as their covariation, while the dynamical model solves the full splicing kinetics. We used the dynamical model throughout our work.

In the following plots, we abbreviate scVelo as “scv”.

***UniTVelo* (Gao et al., 2022)**
UniTVelo employs a top-down, spliced RNA-oriented approach to model transcription dynamics, which directly models spliced RNA profiles using a flexible radial basis function (RBF). It also estimates a unified latent time across all genes by updating cell times along their differentiation path.

In the following plots, we abbreviate UniTVelo as “utv”.

***scTour* (Li, 2023)**
ScTour is a deep learning method that incorporates a variational autoencoder (VAE) and a neural ODE to infer transcriptomic dynamic processes and cell developmental trajectories. In particular, it directly works with total mRNA counts, without relying on the dynamics of spliced and unspliced mRNA. In addition, unlike other methods analyzed in our work, scTour does not output cell-specific velocities on the dimension of each gene. Instead, it projects the estimated velocities to a five-dimensional embedding and visualizes the directions on the low-dimensional field. To compare these results with other methods, we recover the high-dimensional velocity vectors through a decoder and proceed with the replication and local coherence calculations. We describe this in detail in Supplementary Section S3.2.

In the following plots, we abbreviate scTour as “sct”.

***VeloVI* (Gayoso et al., 2024)**
VeloVI is a deep generative method modeling the spliced and unspliced dynamics. It estimates kinetic parameters and latent times for each cell and gene independently, providing an empirical posterior distribution of velocity vectors. The availability of the posterior distribution enables resampling and quantification of the uncertainty in the estimates.

Additionally, the standard VeloVI pipeline includes a preprocessing function that filters out genes with poor linear regression R-squared performance in a steady-state model (i.e., "deterministic" mode, with $R^2 < 0$ or estimated degradation rates $\gamma < 0$) and scales the data. This preprocessing function has the intent of removing "spurious" genes based on the criteria that it deviates too much from a steady-state RNA velocity model. This preprocessing is done via the `velovi.preprocess_data()` function. In our analysis, we tried both versions of running this preprocessing function and skipping this step. We refer to these pipelines as the standard VeloVI and VeloVI without preprocessing, respectively.

In the following plots, we abbreviate standard VeloVI as "velovi" and VeloVI without preprocessing as "velovi_woprep."

#### S1.2. Pseudocode of replicate coherence

The following is the pseudocode for our replicate coherence framework.

```
Input: AnnData object adata with layers "spliced" and "unspliced"

S := adata.layers["spliced"]          # Ncells × Ngenes spliced counts
U := adata.layers["unspliced"]        # Ncells × Ngenes unspliced counts

# Step 1: gene-wise overdispersion estimation (separately for spliced
and unspliced)
for gene in genes:
    theta_S[gene] := estimate_overdispersion( S[:, gene] )    # spliced
overdispersion
    theta_U[gene] := estimate_overdispersion( U[:, gene] )    # unspliced
overdispersion

# Step 2: negative binomial count splitting using the estimated
overdispersions
(S1, S2) := NB_count_split( S, theta_S )    # Split 1 and Split 2 for
spliced matrix
(U1, U2) := NB_count_split( U, theta_U )    # Split 1 and Split 2 for
unspliced matrix

# Construct split AnnData objects (same obs/var as adata, but swapped-in
layers)
split1 := make_adata_like(adata)
split1.layers["spliced"]   := S1
split1.layers["unspliced"] := U1

split2 := make_adata_like(adata)
split2.layers["spliced"]   := S2
split2.layers["unspliced"] := U2

# Step 3: run RNA velocity separately on Split 1 and Split 2
velo1 := run_velocity_method(split1)    # velocity vectors, one per cell
```

```
velo2 := run_velocity_method(split2)

# Step 4: compute replicate coherence per cell
for cell_i in cells:
    replicate_coherence[cell_i] := cosine_similarity( velo1[cell_i, :],
velo2[cell_i, :] )

# Step 5: output
Output: replicate_coherence
```

In this pseudocode: `estimate_overdispersion` refers to our estimation procedure to estimate the overdispersion for each gene based on the count data; `NB_count_split` refers to our count-splitting procedure; `make_adata_like` refers to creating a new single-cell dataset inheriting the same set of cells and metadata; `run_velocity_method` refers to estimating the RNA velocity using a particular method; and `cosine_similarity` refers to computing the cosine similarity between two velocity vectors, which is the replicate coherence.

#### S1.3. Pseudocode for selecting non-informative genes

The following pseudocode shows how we select non-informative genes based on a set of informative genes.

```
Input:
  - genes_mark: Marker genes (GPC genes)
  - AnnData object with spliced/unspliced counts
  - n_sample_scale: scaling factor (n_sample_scale=2 here)
  - bs: number of bins per axis (bs=4 here)

# Step 1: For each gene, compute 3 matching features
Compute:
   (a) fraction of nonzero spliced counts: fnzS[g] := mean(
adata.layers["spliced"][:, g] > 0 )    for each gene g
   (b) fraction of nonzero unspliced counts: fnzU[g] := mean(
adata.layers["unspliced"][:, g] > 0 ) for each gene g
   (c) RNA velocity R-squared based on scVelo deterministic mode

# Step 2: Discretize the 3D feature space into bins and assign each
marker and control gene to its nearest bin.
   df_mark := rows for genes in genes_mark, with columns (fnzS, fnzU, r2)
   df_ctrl := rows for genes not in genes_mark, with columns (fnzS, fnzU,
r2)
   # Create a 3D grid of bin centers using marker ranges
   bins1 := Partition equal-length bins from min(df_mark.fnzS) to
max(df_mark.fnzS)
   bins2 := Partition equal-length bins from min(df_mark.fnzU) to
max(df_mark.fnzU)
```

```
   bins3 := Partition equal-length bins from min(df_mark.r2) to
max(df_mark.r2)
   bin_centers := all triples (b1, b2, b3) from bins1×bins2×bins3 # total
bins = bs^3

   # Assign each gene to its nearest bin center (kNN with k=1)
   membership_mark[g] := argmin_over_bins Euclidean distance between
feature(g) and bin_centers[bin] for g in df_mark
   membership_ctrl[g] := argmin_over_bins Euclidean distance between
feature(g) and bin_centers[bin] for g in df_ctrl

   # Compute target counts per bin from markers
   freq_mark[bin] := number of genes in genes_mark assigned to each bin

   # Compute bin-to-bin distances for neighbor fallback
   bin_dist[bin_i, bin_j] := Euclidean distance between
bin_centers[bin_i] and bin_centers[bin_j]

   For each seed of count splitting:
     genes_select := empty set
     For each bin:
       target := freq_mark[bin] * n_sample_scale
       # Prefer control genes from the same bin
       add up to target unused control genes with membership_ctrl = bin
       # If still short, fill the remainder using unused control genes
from the closest bin in (fnzS, fnzU, r2) space
       while N_selected_from_bin < target:
         Choose the nearest bin_nbr (not yet used for this bin)
         Add as many unused control genes to genes_select from bin_nbr as
needed

# Step 3: Use the selected genes to construct matched control AnnData
objects for the full dataset and for each split-replicate dataset.
   adata_ctrl_total := adata_full[:,genes_selected]
   adata_ctrl_split1 := adata_split1[:,genes_selected]
   adata_ctrl_split2 := adata_split2[:,genes_selected]
```

#### S1.4. Discussion about count splitting compared to other potential strategies

We note that during revision, a reviewer suggested an alternative to count splitting based on subsampling cells. Under this approach, cells that appear in multiple subsamples can be compared across fits using cosine similarity. The idea is straightforward to implement and, with careful execution, could provide a stability check for RNA-velocity estimates. In practice, however, it might be difficult to control cell-type composition across subsamples. Poorly balanced subsampling could substantially alter the neighborhood graph, making downstream velocity estimates difficult to compare across runs. Moreover, because subsampling could reduce the number of cells per fit, one may end up with too few cells for reliable RNA-velocity estimation and/or too small an overlap between subsamples for meaningful comparisons. In contrast, count splitting preserves the full sample size and yields paired splits, ensuring sufficient data for fitting RNA-velocity methods and for direct comparisons between splits.

In addition, because comparisons rely on overlapping cells across subsamples, the resulting datasets might not be mutually exclusive. This dependence complicates the interpretation of velocity-vector comparisons, since the cosine similarity might reflect shared signal from overlapping cells rather than independent variability. Our proposed procedure incorporating count splitting avoids this issue by producing independent splits, which makes cosine-similarity comparisons easier to interpret as capturing variability induced by the split rather than shared cells across subsamples.

### S2. Overview of datasets

We provide summary statistics of the mouse gastrulation [Pijuan-Sala et al., 2019] ("erythroid"), the pancreatic endocrinogenesis development [Bastidas-Ponce et al., 2019] ("pancreas"), and the human cerebral cortex development [Trevino et al., 2021] ("brain") datasets.

*Table S1: Summary of each dataset*

| | erythroid | pancreas | pancreas without pre-endocrine | brain |
|---|---|---|---|---|
| Number of cells | 9815 | 3696 | 3104 | 38,262 |
| Number of genes (before selecting highly-variable genes) | 53,801 | 27,998 | 27,998 | 32,648 |
| Average non-zero spliced counts across cells | 4.649 | 2.653 | 2.715 | 2.228 |
| Fraction of genes with non-zero spliced expression | 38.86% | 63.40% | 62.80% | 75.06% |

In the erythroid dataset, the cell types were labeled based on their expression of marker genes *Ptprc* (encoding CD45), *Kit*, *Csf1r*, and *Fcgr3* (encoding CD16). These markers were examined within clusters obtained from a shared nearest-neighbor graph and by applying the Louvain algorithm. Cells that did not exhibit a unique marker profile were merged with closely related clusters, ensuring that the annotations were reliable and supported by established gene signatures.

- **Data version/source:** The data is installed along with the scVelo package (https://pypi.org/project/scvelo/0.3.1/). See https://scvelo.readthedocs.io/en/stable/scvelo.datasets.gastrulation_erythroid.html#scvelo.datasets.gastrulation_erythroid for documentation on the dataset. The celltype label is stored under `.obs[celltype]` in the anndata object.

In the pancreas dataset, a graph-based clustering approach was used, yielding clusters of distinct pancreatic lineages at various embryonic stages. The clusters were labeled based on the expression of well-studied marker genes functionally related to pancreas development, including *Dlk1* (MPCs), *Cpa1* and *Myc* (tip), *Notch2* (trunk), *Ptf1a*, *Cpa1*, *Cel*, *Rbpjl* (acinar), *Sox9*, *Anxa2* and *Bicc1* (ductal), *Ngn3* and *Hes6* (EPs), *Fev*, *Cck* and *Neurod1* (*Fev* high), and *Rbp4*, *Pyy* and *Chgb* (endocrine).

- **Data version/source:** The data is installed along with the scVelo package (https://pypi.org/project/scvelo/0.3.1/). See https://scvelo.readthedocs.io/en/stable/scvelo.datasets.pancreas.html#scvelo.datasets.pancreas for documentation on the dataset. The celltype label is stored under `.obs['clusters']` in the anndata object.

In the brain dataset, cells were annotated through a combination of unsupervised clustering and manual curation using known marker genes. After clustering both scRNA-seq and scATAC-seq profiles, the authors assigned cell type identities based on the expression and gene activity scores of established lineage markers from previous studies. Annotations were cross-validated using external human cortical scRNA-seq datasets and canonical correlation analysis (CCA) to integrate chromatin and transcriptomic data. Additionally, gene regulatory links between enhancers and genes were established based on co-variation, reinforcing the distinct molecular identity of each annotated cluster.

- **Data version/source:** The data was downloaded from https://github.com/GreenleafLab/brainchromatin, where the URLs of the relevant files are located in links.txt file (i.e., https://github.com/GreenleafLab/brainchromatin/blob/main/links.txt). Specifically, we use the files:
    - https://atrev.s3.amazonaws.com/brainchromatin/multiome_spliced_rna_counts.tsv.gz
    - https://atrev.s3.amazonaws.com/brainchromatin/multiome_unspliced_rna_counts.tsv.gz
    - https://atrev.s3.amazonaws.com/brainchromatin/multiome_cell_metadata.txt
    - https://atrev.s3.amazonaws.com/brainchromatin/multiome_cluster_names.txt

The version for this repository (Commit f8f509b) is linked to the original publication (https://www.sciencedirect.com/science/article/pii/S0092867421009429). The celltype label is stored under `.obs[cluster_name]` in the anndata object.

### S3. Additional details about our method and analyses

#### S3.1. Additional details on how each dataset was processed

**Procedure to estimate overdispersions**
We apply the Gamma-Poisson generalized linear model (GLM) on each spliced and unspliced dataset separately to estimate the overdispersion parameters, using the function `glm_gp` in the R package `glmGamPoi`. By default, `glm_gp` accounts for sample-specific differences in library size (e.g., sequencing depth) via the `size_factor` parameter.

Specifically, we do not adjust for covariates during this overdispersion estimation procedure, since the datasets we analyze in the paper do not include clear and available covariates that are needed to be adjusted for. Moreover, the prescribed framework in Neufeld et al. (2023) does not describe a count splitting procedure that accounts for covariates. Even though `glmGamPoi` can estimate the overdispersion with covariates, it is not immediately clear how to incorporate those covariate-adjusted estimates into our count-splitting workflow.

**scVelo**
First, we filter and normalize highly variable genes, then compute moments. Then, PCA, nearest neighbor graph, and UMAP are computed in order. With all these ingredients, we recover the gene expression dynamics and compute the velocity vectors. For erythroid data, we perform a batch correction using bbknn [Polański et al., 2020] immediately after the moments are computed and before applying PCA.

**UniTVelo**
We start by configuring the model environment and then run the model fitting function. A UMAP is then computed, where the erythroid dataset undergoes batch correction, whereas the pancreas and brain datasets do not.

**scTour**
We follow the scTour pipeline and start by calculating the quality control metric. Then we identify highly variable genes and feed the data object into scTour Trainer. The low-dimensional vector field is computed, which we later use to recover the high-dimensional velocity vectors and the associated velocity graph (details illustrated in a later section). Note that scTour does not compute the UMAP. When we need the UMAP for visualization, we follow a procedure that involves computing moments, performing PCA, identifying neighbors, and then using UMAP. For the erythroid data, we do a batch correction between the moment and PCA computation.

**VeloVI**
First, we filter and normalize highly variable genes, then compute moments. The standard VeloVI runs a preprocess_data function at this stage, further filtering out poorly performing genes. The VeloVI without preprocessing skips this step. The object is then fed into the VELOVI

trainer. The velocity estimates are finally obtained by the function *add_velovi_outputs_to_adata*. Like scTour, VeloVI also does not compute a UMAP by default. We again follow the procedure of computing moments, performing PCA, identifying neighbors, and using UMAP. For the erythroid data, we do a batch correction between the moment and PCA computation.

#### S3.2. Additional details on how to compute replicate coherence for scTour

In RNA velocity computation, scTour differs from other methods in the type of information utilized. While scVelo, UniTVelo, and VeloVI leverage spliced and unspliced RNA dynamics to directly compute velocities, scTour bypasses this distinction and produces a low-dimensional vector field. To make scTour comparable to other methods, we manually decode its five-dimensional vectors to recover high-dimensional velocities. This is important since our replicate coherence relies on both splits having the same “features”. Abstractly, training scTour twice, once on each split, might result in two “unaligned” five-dimensional latent spaces. Hence, our following procedure reconstructs the velocity vector with dimensionality equal to the number of genes. In this way, we can compute the correlation of scTour’s RNA velocity vector across the two splits meaningfully.

This decoding requires specifying a timestep parameter, where a larger timestep predicts a velocity vector further from the current state. To mitigate the impact of different parameter choices, we recover velocities at ten evenly spaced timesteps ranging from 0.02 to 0.2 and compute their average. Additionally, we address potential pseudotime reversal in the decoded results by verifying the correlation between scTour pseudotime and the recovered pseudotime.

#### S3.3. Gene correlation in the erythroid dataset

The histograms show the correlation of the genes between two splits, for spliced and unspliced counts separately, after running count splitting on the mouse erythroid dataset. The correlations are mostly around 0, indicating that count splitting effectively creates independent subsets that can reasonably be considered technical replicates.

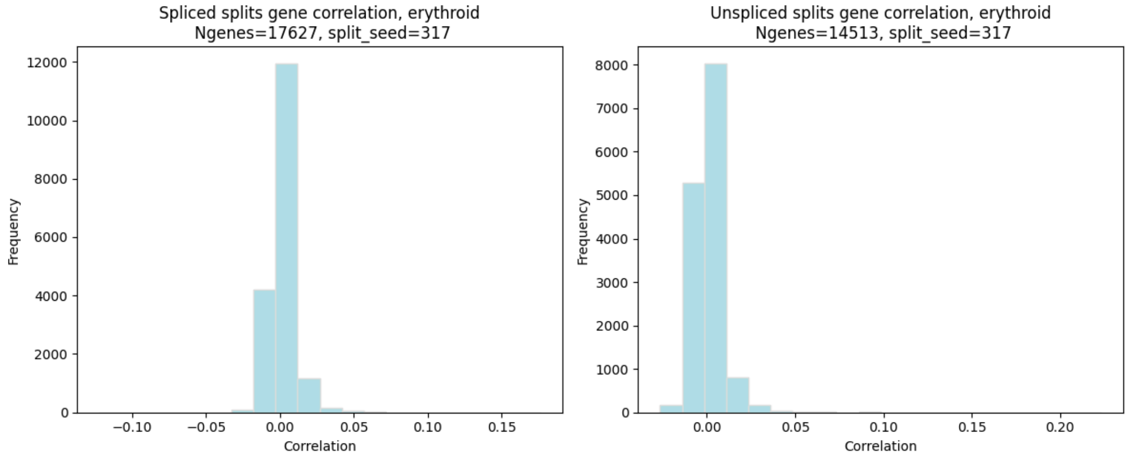

*Figure S1: Plots showing the correlation of genes between two splits for the erythroid dataset. The left and right columns denote plots based on the spliced and unspliced counts.*

**DESeq2-like overdispersion estimates:**
The primary objective of the following investigation is to demonstrate that our count splitting framework is robust to the specific method used for overdispersion estimation. While our standard workflow utilizes a gene-specific Gamma-Poisson GLM, we empirically test whether alternative estimation strategies, such as DESeq2 which shares information across genes, materially affect the independence properties of the splits.

Although DESeq2 provides a robust framework for information-sharing, it was originally optimized for bulk RNA-seq data. To maintain precise control over the smoothing process in a single-cell context, we re-implement a DESeq2-like approach that entails:

- Initial Estimates: Fitting gene-specific overdispersion parameters to establish a baseline mean-variance relationship between the overdispersion and the sequencing depth.
- Trend Fitting: Fitting a global trend line (as a function of the mean expression) across all genes to capture the shared dispersion structure.
- Shrinkage: Shrinking gene-specific estimates toward this global trend to reduce the impact of cell-specific noise being misattributed as overdispersion. This implementation confirms that substantial smoothing of dispersion information does not meaningfully alter the final RNA velocity stability results, reinforcing the validity of our original gene-specific approach.

To illustrate the impact of this overdispersion shrinkage, we purposefully increase the smoothing above what DESeq2 typically does. We plot the shrunk vs. original overdispersion, along with the trendline regressing the overdispersion onto the sequencing depth. We notice that despite the aggressive smoothing, many genes did not substantially change in their overdispersion.

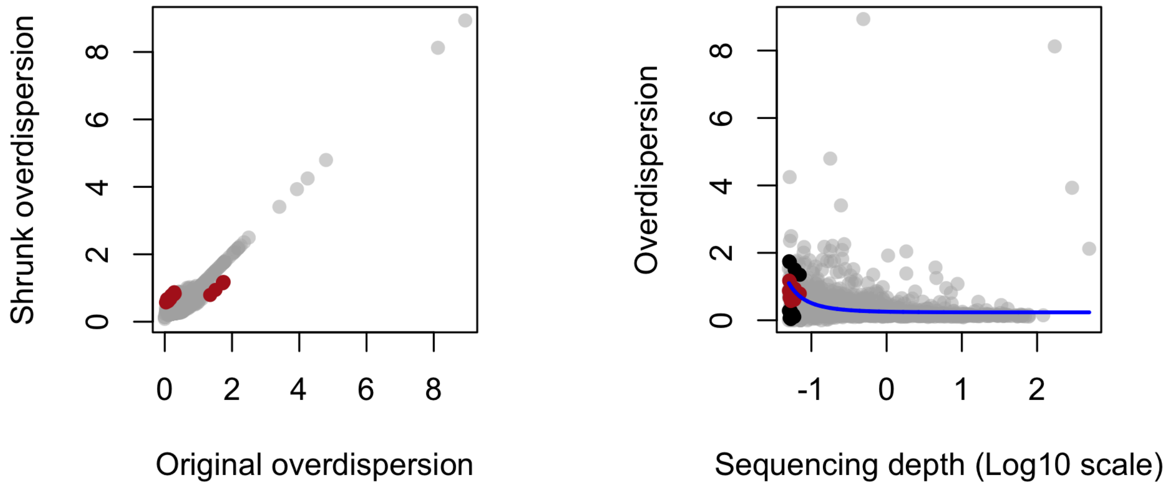


*Figure S2: (Left) Shrunk overdispersion versus original overdispersion, when applied to the spliced counts on the erythroid dataset. (Right) The overdispersion versus the sequencing depth (log-10 scale). In both plots, each point denotes a gene, and the genes with the most substantial*

*changes in overdispersion are marked in red (overdispersion changed by more than 0.5; others: gray). The blue line denotes the trendline, and the black point denotes the original overdispersion before shrinkage.*

Using these alternative overdispersion estimates, we perform count splitting and then compute the correlation between gene expression counts across the two splits, separately for spliced and unspliced genes. Similar to the previous results, the correlations are centered at 0, indicating independence between gene expression in the two splits.

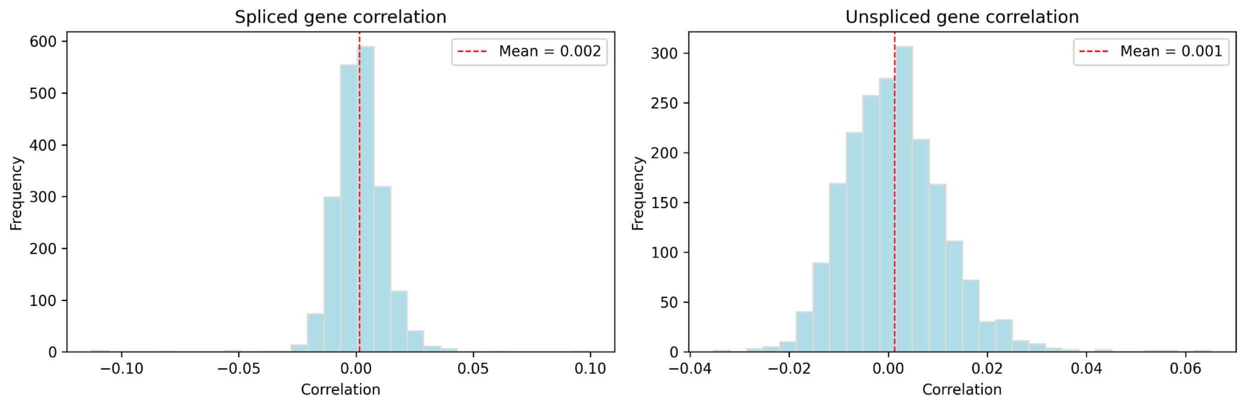


*Figure S3: Plots showing the correlation of genes between two splits for the erythroid dataset using the DESeq2-like overdispersion estimates. The left and right columns denote plots based on the spliced and unspliced counts.*

In addition, we compute the replicate coherence from scVelo's results. The median and mean replicate coherence values are close to the ones obtained from the original overdispersion estimates.

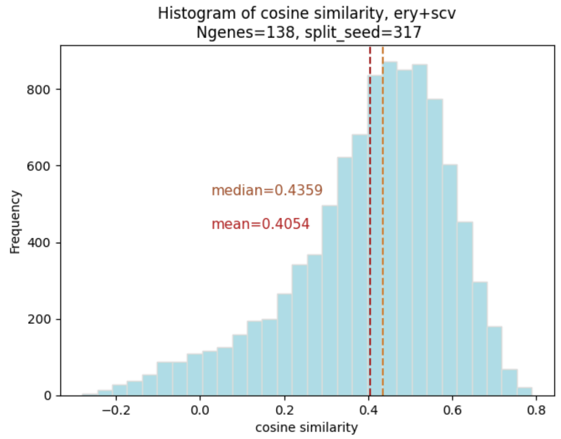


*Figure S4: Histogram of replicate coherence for erythroid using the DESeq2-like overdispersion estimates and scVelo. This is to be compared against Figure 2b for "scVelo".*

Our results indicate that the "replicate coherence" and the independence between technical replicates remain stable across different overdispersion models, suggesting that the framework reliably preserves the underlying biological signal even when the overdispersion parameters are estimated through more complex pooling methods.

We note that there are many ways to estimate the overdispersion parameter due to its intrinsic statistical difficulty. Notably, when the data becomes very sparse, the log-likelihood function of the overdispersion parameter becomes very flat, meaning that the overdispersion magnitude can differ a lot with little change to the log-likelihood. This means that while glmGamPoi is the implementation that we have found successful in our data analysis both in computational efficacy and statistical accuracy, other overdispersion estimation procedures can be considered on a case-by-case basis.

#### S3.4. Overdispersion vs. correlation vs. sparsity

The plots below show the gene correlations between splits, estimated overdispersion parameters, and sparsity of the erythroid data. Here, we threshold the overdispersion to be at most 100 for visual clarity (along the x-axis), even if the estimated overdispersion is much higher than 100.

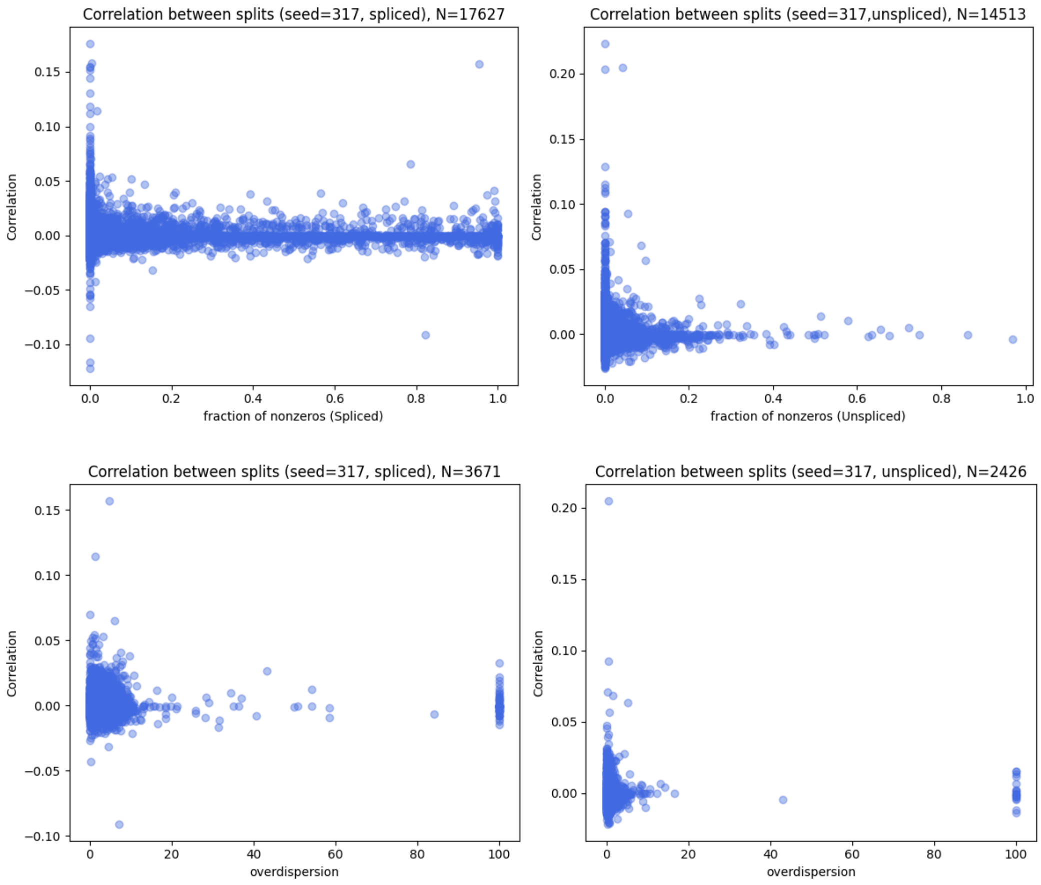

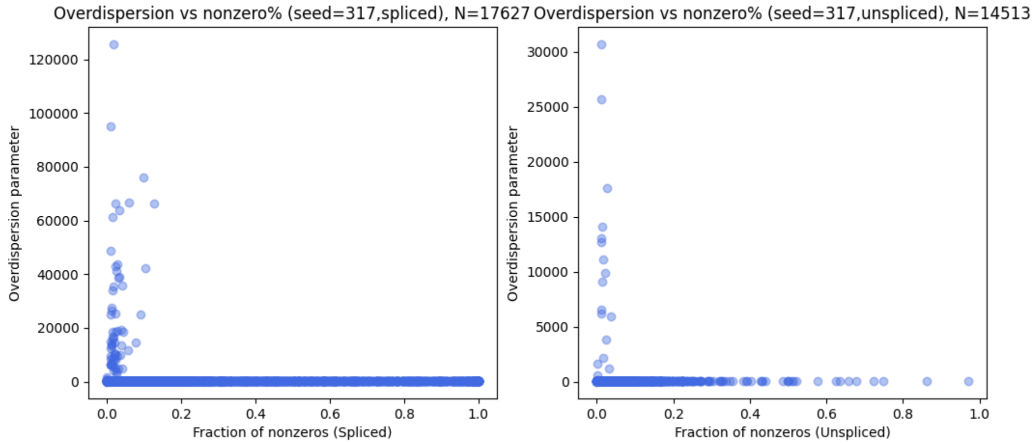


*Figure S5: Plots showing the relationship between the correlation between the two splits (via negative-binomial count splitting), the estimated overdispersion parameter for that gene (where a higher overdispersion denotes a gene that has more like a Poisson), and the fraction of cells with a non-zero sequencing count for that gene, where each point is a gene. This plot is made for the erythroid dataset. The left and right columns denote plots based on the spliced and unspliced counts.*

#### S3.5. Table of intersection and union of genes used in velocity

The tables in this section show the size of the intersection and union of genes used to compute velocity vectors in two splits for each method on each dataset. Note that we used the union of genes in the main text to compute replicate coherence, with genes appearing only one split padded with 0's. However, the results did not vary largely whether we used the union or the intersection in practice.

*Table S2: Number of genes in the intersection or union between the two splits after applying each RNA velocity method, where the gene sets denote which genes that method used, both based on the gene being a highly-variable gene in that split, and also whether the gene is used in the RNA velocity vector*

Erythroid

| | scVelo | UniTVelo | scTour | VeloVI (std) | VeloVI (w/o prep) |
|---|---|---|---|---|---|
| intersection | 212 | 1480 | 1215 | 89 | 1480 |
| union | 417 | 2520 | 2785 | 167 | 2520 |

Pancreas

| | scVelo | UniTVelo | scTour | VeloVI (std) | VeloVI (w/o prep) |
|---|---|---|---|---|---|
| intersection | 716 | 1487 | 1486 | 506 | 1487 |
| union | 1102 | 2513 | 2514 | 743 | 2513 |

Pancreas without pre-endocrine

| | scVelo | UniTVelo | scTour | VeloVI (std) | VeloVI (w/o prep) |
|---|---|---|---|---|---|
| intersection | 751 | 1497 | 1452 | 545 | 1497 |
| union | 1161 | 2503 | 2548 | 777 | 2503 |

Brain (results using informative GPC genes)

| | scVelo | UniTVelo | scTour | VeloVI (std) | VeloVI (w/o prep) |
|---|---|---|---|---|---|
| intersection | 67 | 144 | 184 | 65 | 144 |
| union | 77 | 150 | 184 | 78 | 150 |

We note the following observation regarding the standard VeloVI and VeloVI withoout preprocessing: VeloVI's preprocessing via the `velovi.preprocess_data()` function filters genes via a poor linear regression R-squared performance in a steady-state model (i.e., scVelo in "deterministic" mode, with $R^2 < 0$ or estimated degradation rates $\gamma < 0$). We noted this substantially filtered out many genes. Across multiple datasets (Figures 2-4), we observed that VeloVI run without `velovi.preprocess_data()` tends to yield higher replicate coherence than the default VeloVI pipeline. A plausible explanation is that the preprocessing step performs substantial gene filtering based on a steady-state model; in our settings, this can remove a large fraction of genes and thereby reduce the information available for learning RNA velocity via a deep-learning approach. In contrast, scVelo uses a similar number of genes. However, scVelo's RNA velocity modeling uses a simpler parametric model, and may therefore be less sensitive to this type of filtering.

Importantly, this pattern does not appear to be driven by count-splitting-induced sparsity or by deep learning per se, since VeloVI without preprocessing remains stable in our experiments. We do not advocate against gene filtering in general; rather, this result illustrates the intended use of replicate coherence as an additional diagnostic to help practitioners assess how sensitive a fitted velocity field is to pipeline choices, including preprocessing.

#### S3.6. UMAP: Erythroid between two splits

The plots in this section show the velocity estimates of splits plotted on UMAPs for each method on the erythroid dataset. Note that these UMAPs are recovered based on counts in each split rather than the embedding from the original erythroid data object.

Note that since each split receives roughly half the counts (while being independent), the UMAP coordinates based on Split 1 do not necessarily resemble the UMAP based on Split 2. However, the utility of the following plots is to visually diagnose the source of potential low replicate coherence based on the RNA velocity field.

#### scVelo (deterministic mode)

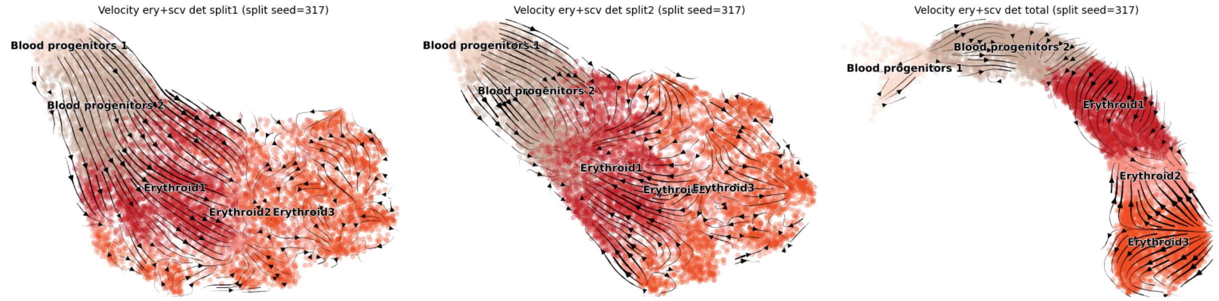


#### scVelo (dynamical mode)

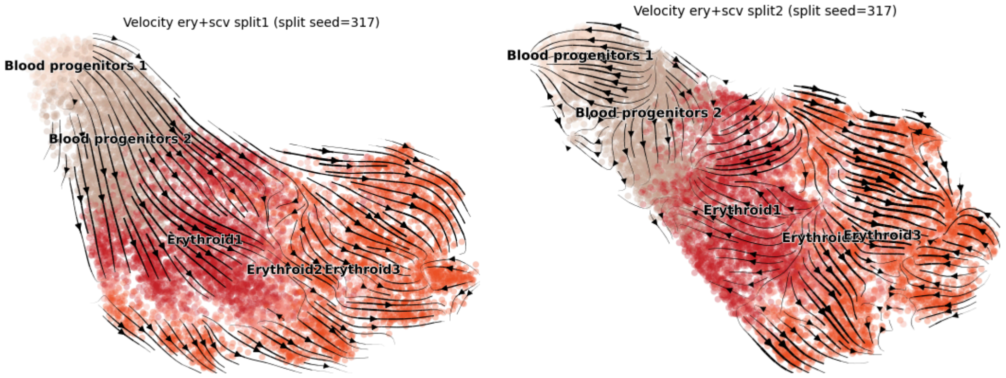


#### UniTVelo

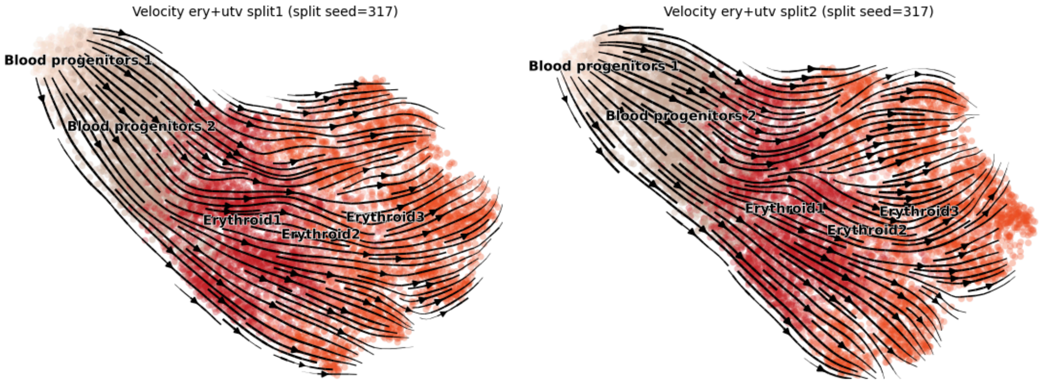


#### scTour

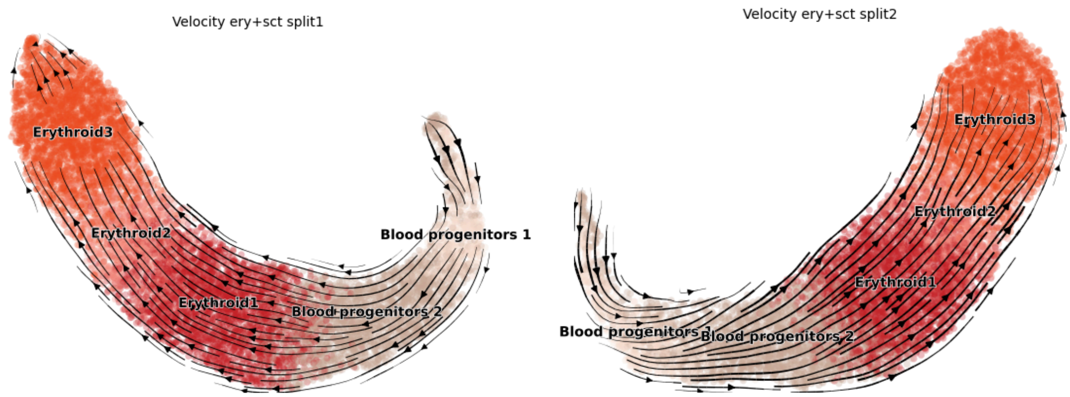


#### VeloVI (standard)

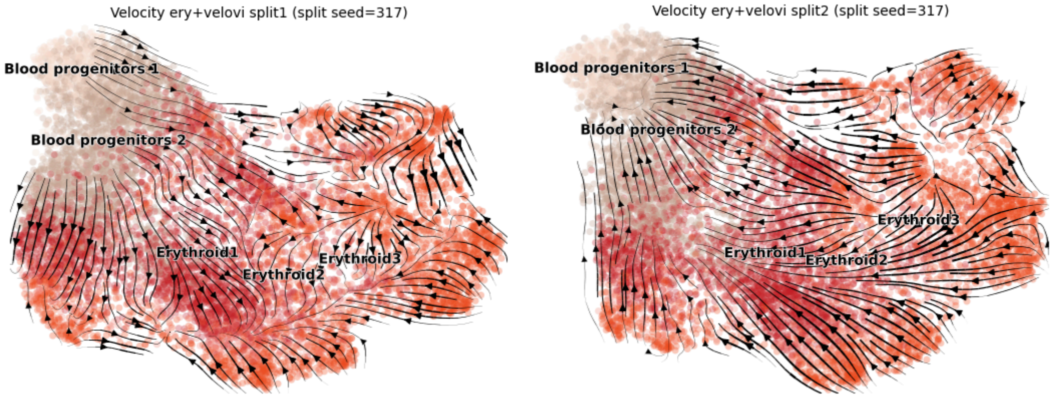


#### VeloVI without preprocessing

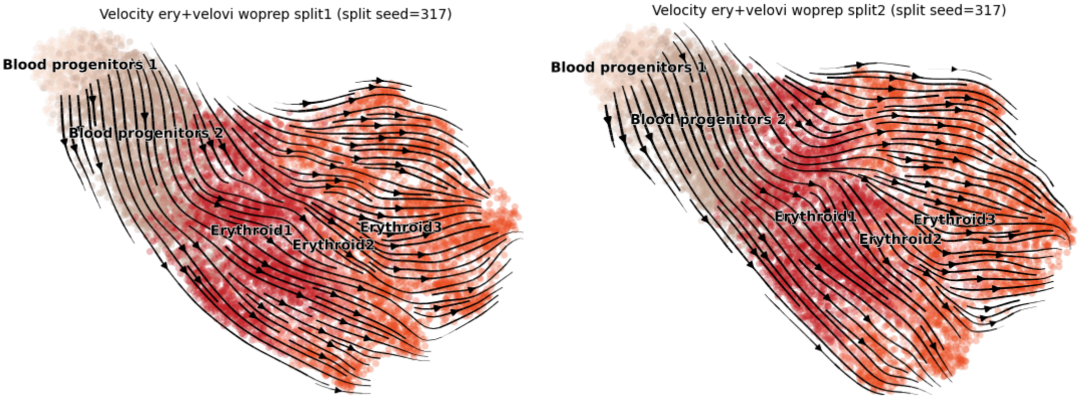


*Figure S6: UMAP showing the RNA velocity flow, based on applying each RNA velocity method on the two splits of the erythroid data after applying negative-binomial count splitting. The cell type colors are the same as in Figure 2. The exception is for scVelo (deterministic), where we include 3 plots where the right-most plot is of the total counts, resembling the plots in Figure 2a in the main text.*

### S3.7. VeloVI extrinsic and intrinsic uncertainty on pancreas

VeloVI provides two measures of uncertainty: intrinsic and extrinsic. Intrinsic uncertainty quantifies the variance in cosine similarity among samples of the velocity vector. In contrast, extrinsic uncertainty captures the variance in cosine similarity between velocity samples and future cell states, both of which are calculated with respect to their respective means. The figures below display the intrinsic and extrinsic uncertainty, at a log scale, for both standard VeloVI and VeloVI without preprocessing applied to the pancreas dataset. In the plots, higher uncertainty values are represented by redder colors, while lower uncertainty values are depicted in shades of blue.

Note that extrinsic uncertainty values from both methods have a more negative range compared to intrinsic uncertainty values. This suggests lower uncertainty or, equivalently, smaller variances in cosine similarity when comparing velocity samples with future cell states, as opposed to variances within the sampled velocity vectors.

**VeloVI (standard)**

Extrinsic uncertainty

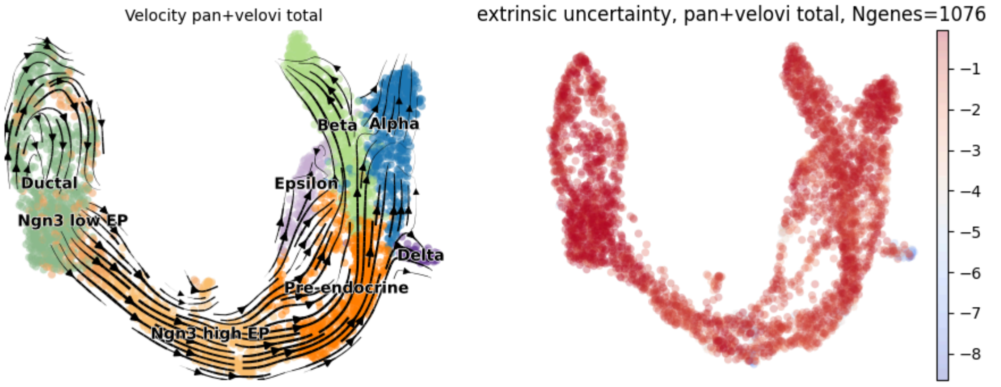


Intrinsic uncertainty

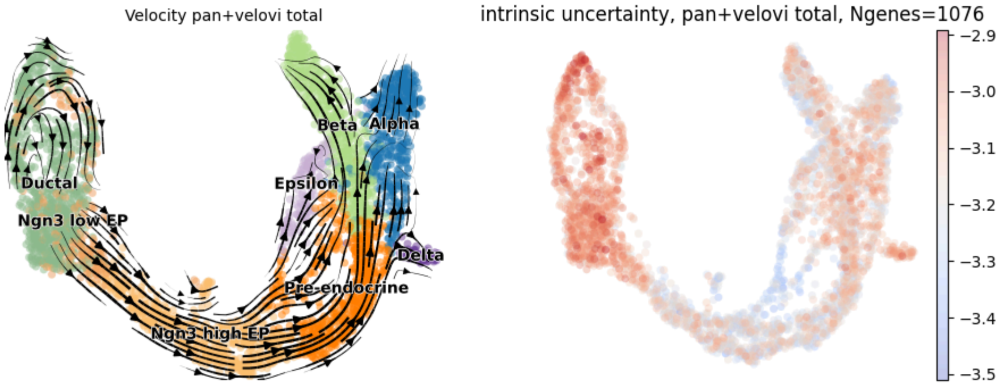

*Figure S7: The extrinsic and intrinsic uncertainty computed via standard VeloVI on the pancreas dataset. The UMAP with RNA velocity flow on the left is identical to that shown in Figure 3a. The plots on the right denote the uncertainty, where red indicates higher uncertainty (i.e., the log of the variance is large), while blue represents low uncertainty (i.e., the log of the variance is small). The color bar in each figure represents the estimated variance of the RNA velocity correlation based on VeloVI, displayed on a log scale. The cell type colors are the same as in Figure 3.*

**VeloVI without the preprocess step**

Extrinsic uncertainty

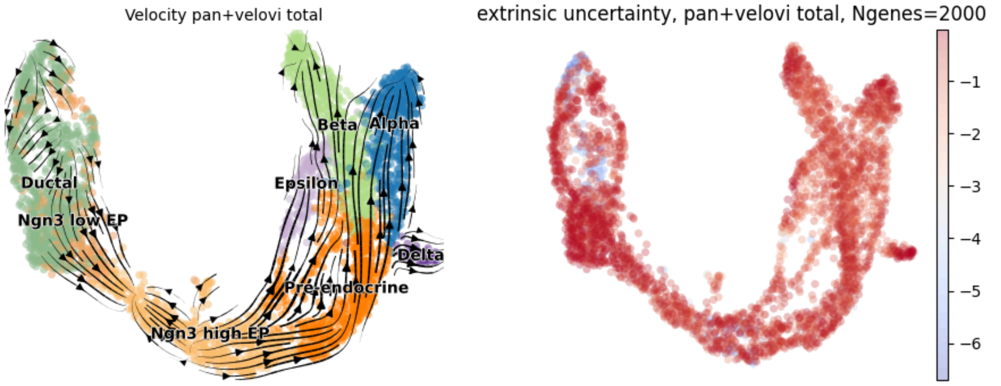


Intrinsic uncertainty

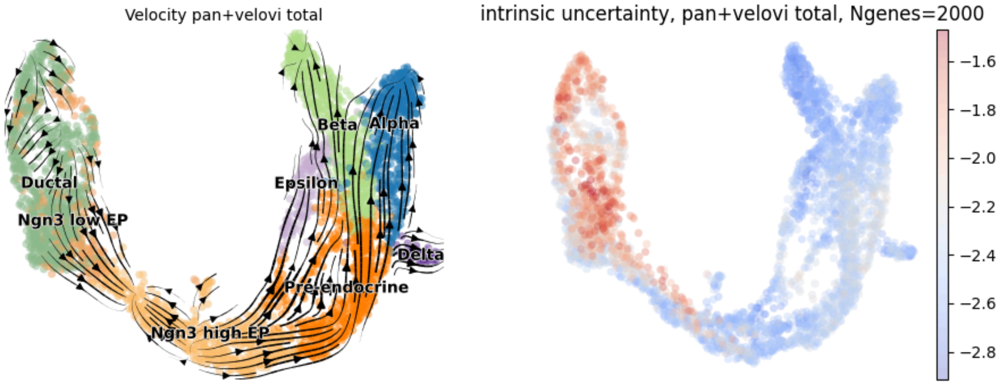


*Figure S8: The same layout as Supplementary Figure S7, except using VeloVI without preprocessing.*

#### S3.8. UMAP: Pancreas & pancreas without pre-endocrine 5 methods

Below are the velocity estimates plotted on the UMAP embedding inherited from the original pancreas dataset. The left figure shows the velocity estimated by each method on the full pancreas data, while the right one shows the velocity estimates on the pancreas data without pre-endocrine.

**scVelo**

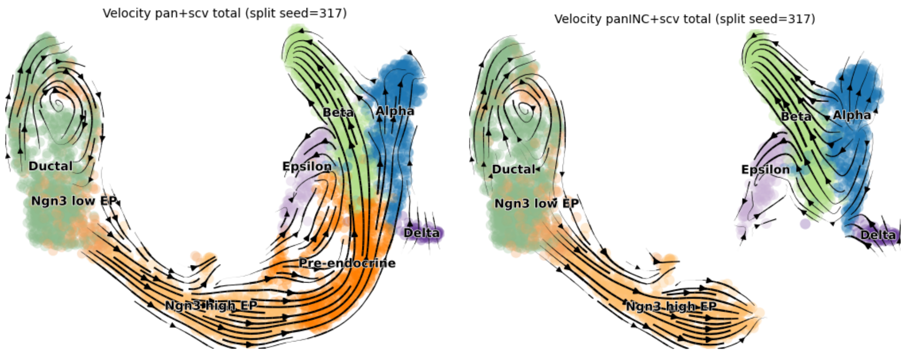


**UniTVelo**

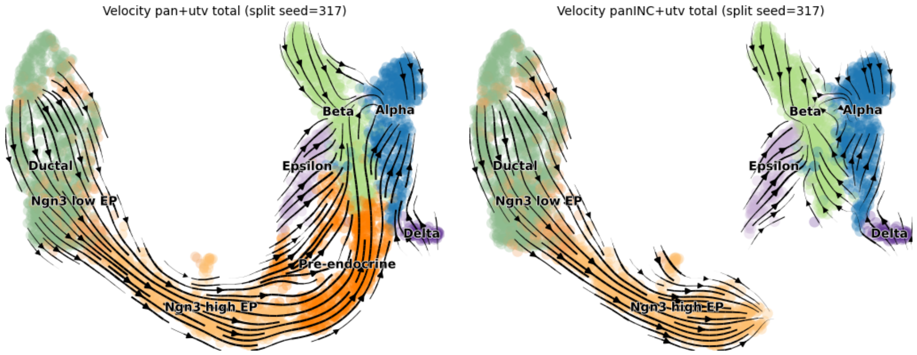


**scTour**

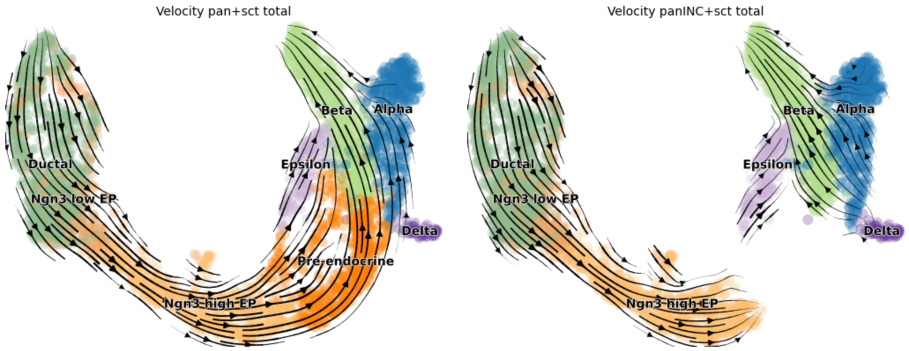


**VeloVI (standard)**

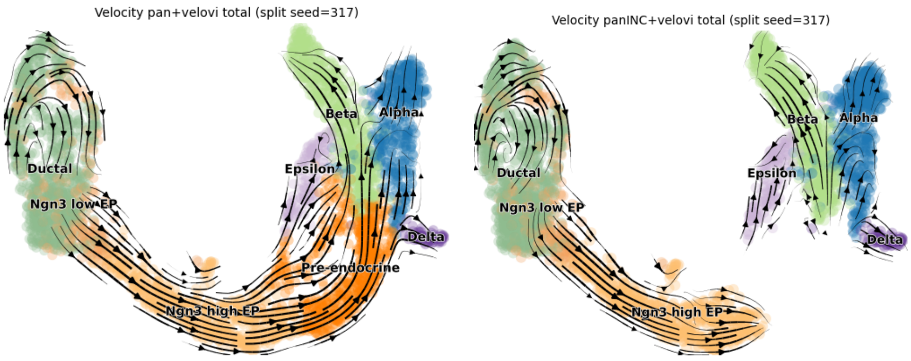


**VeloVI without preprocessing**

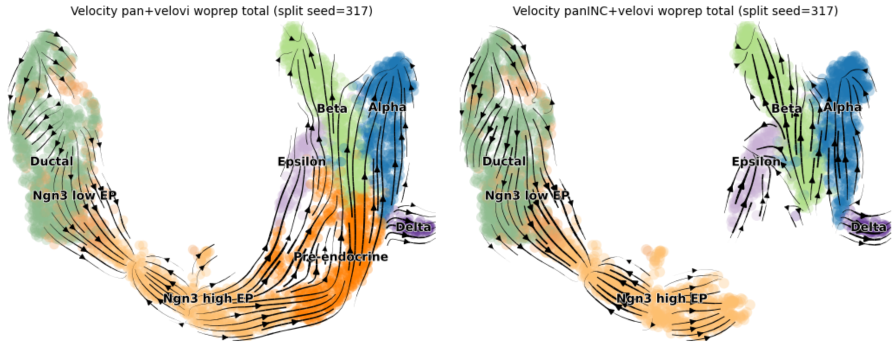


*Figure S9: The UMAPs with RNA velocity flow from each RNA velocity method for the pancreas dataset, with (left column) and without the pre-endocrine cells (right column). The cell type colors are the same as in Figure 3 in the main text.*

#### S3.9. Local/replicate coherence in the datasets

This section presents the summary of local and replicate coherence across five RNA velocity methods for the pancreas, erythroid, and brain datasets. The distributions are shown as boxplots, following the same format as Figure 2c in the main text. Median values for each distribution are provided in tables following each boxplot.

**Erythroid (additional results)**

In addition to the five methods mentioned in the main text, we also run scVelo in deterministic mode, which is equivalent to Velocyto (La Manno, 2018), on the erythroid dataset. The distributions of local and replicate coherence obtained from one seed are shown in the following figure. We note that the replicate coherence for this method is lower than that shown in Figure 2 of the main text.

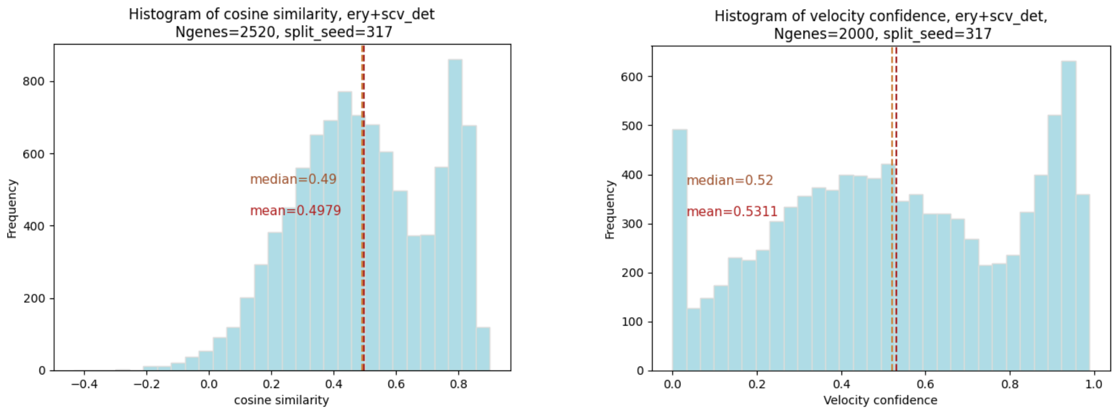


*Figure S10: Histogram of the local and* replicate *coherence of scVelo deterministic mode (Velocyto) for the erythroid dataset.*

#### Pancreas

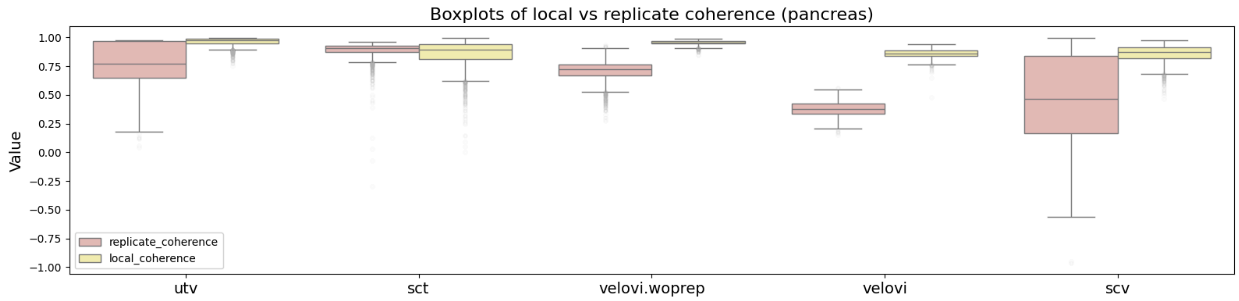


*Figure S11: Boxplot of the local and replicate coherence across the five RNA velocity methods for the pancreas dataset, in the same format as Figure 2b in the main text.*

*Table S3: Median value of replicate or local coherence (among all the cells) for the pancreas dataset across the five RNA velocity methods.*

| | utv | sct | velovi.woprep | velovi | scv |
|---|---|---|---|---|---|
| Replicate | 0.7727 | 0.9063 | 0.7225 | 0.3786 | 0.4658 |
| Local | 0.9787 | 0.8969 | 0.9580 | 0.8634 | 0.8716 |

#### Brain

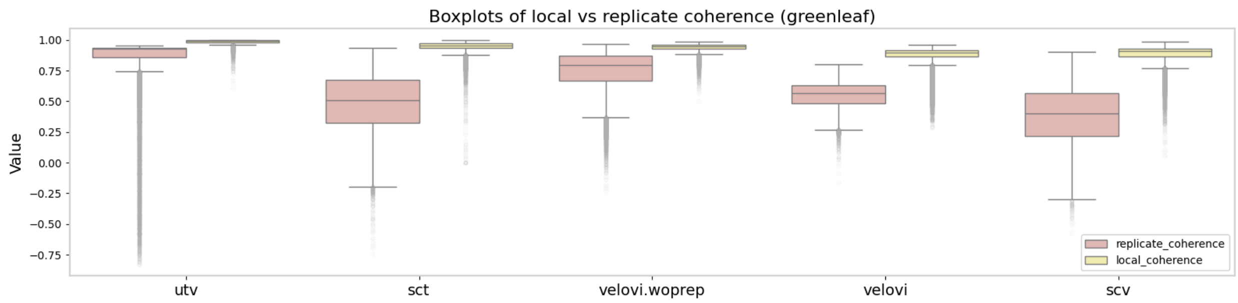


*Figure S12: Boxplot of the local and replicate coherence across the five RNA velocity methods for the brain dataset, in the same format as Figure 2b in the main text.*

*Table S4: Median value of replicate or local coherence (among all the cells) for the brain dataset across the five RNA velocity methods.*

| | utv | sct | velovi.woprep | velovi | scv |
|---|---|---|---|---|---|
| Replicate | 0.9258 | 0.5099 | 0.7949 | 0.5674 | 0.3988 |
| Local | 0.9893 | 0.9554 | 0.9462 | 0.8969 | 0.9082 |

Below are the velocity estimates and UMAP colored by local coherence and replicate coherence, as applied to the two versions of the pancreas data using different methods. The leftmost plot shows the velocity estimates plotted on the UMAP embedding. The middle plot shows the local coherence, and the rightmost plot shows the replicate coherence.

**UMAPs of pancreas data, colored by local and replicate coherence.**

**scVelo**

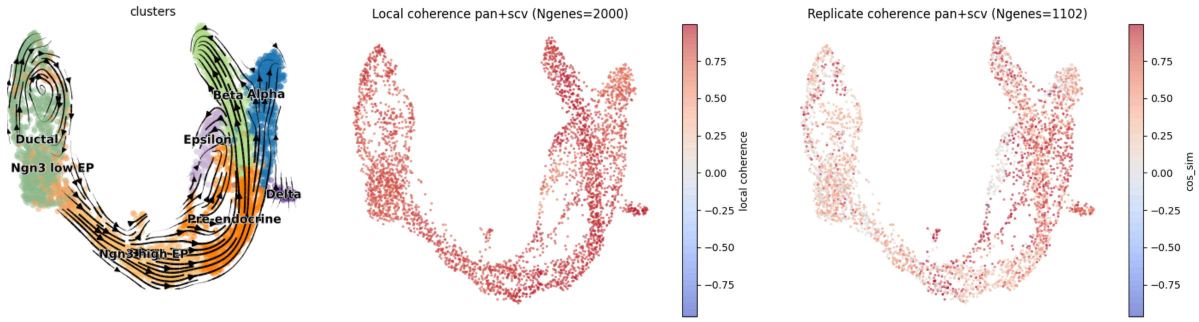


**UniTVelo**

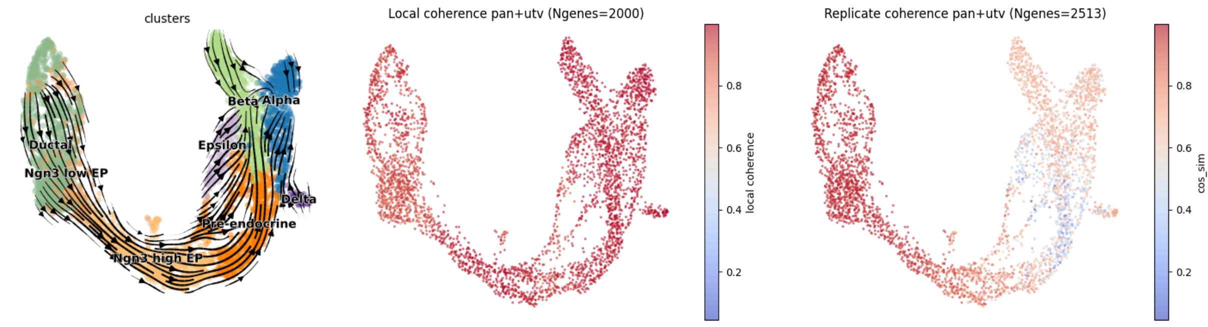


#### scTour

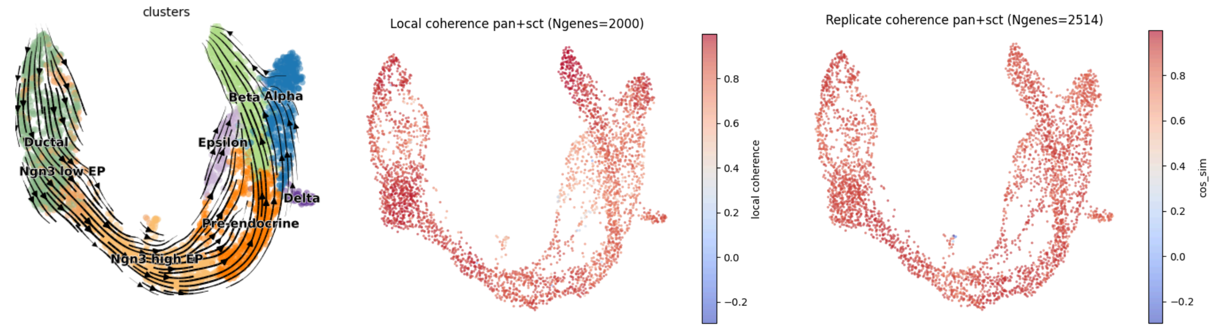


#### VeloVI (standard)

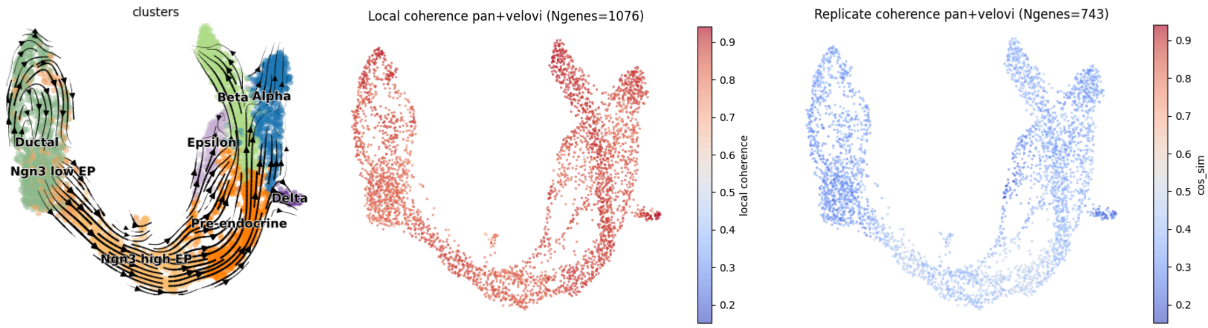


#### VeloVI without preprocessing

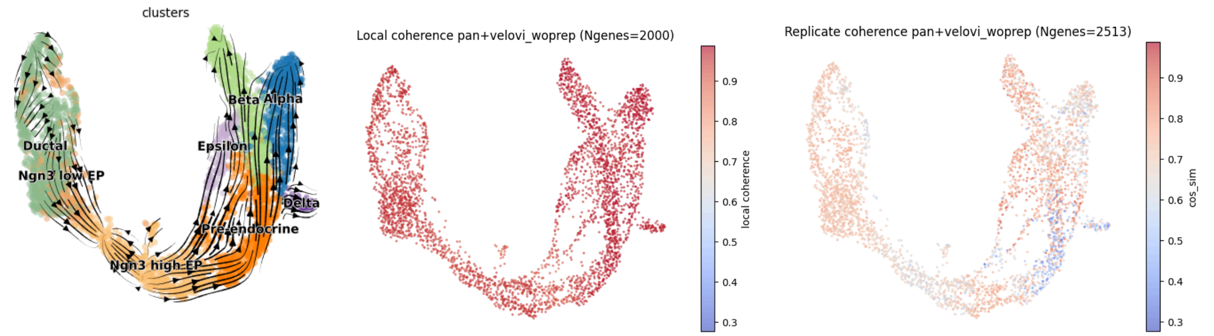


*Figure S13: The comparison of the local coherence (i.e., “velocity confidence”, middle column) against our replicate coherence (i.e., “velocity cosine similarity”, right column) in the pancreas dataset. The UMAPs with RNA velocity flow are shown in the left column as reference, which*

*are the same as those in Supplementary Figure S9. The cell type colors are the same as in Figure 3, and the same color bar is shown for each of the coherence plots, where red denotes high coherence (i.e., low variability) and blue denotes low coherence (i.e., high variability).*

**<u>UMAPs of pancreas without pre-endocrine, colored by local and replicate coherence.</u>**

**scVelo**

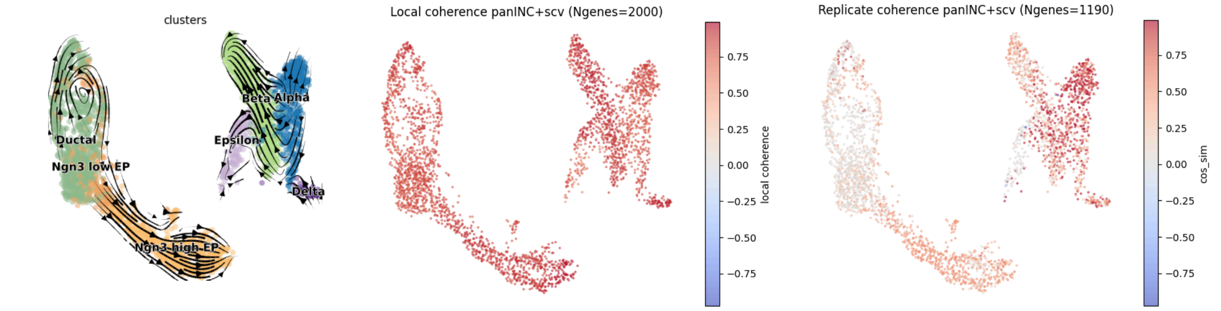


**UniTVelo**

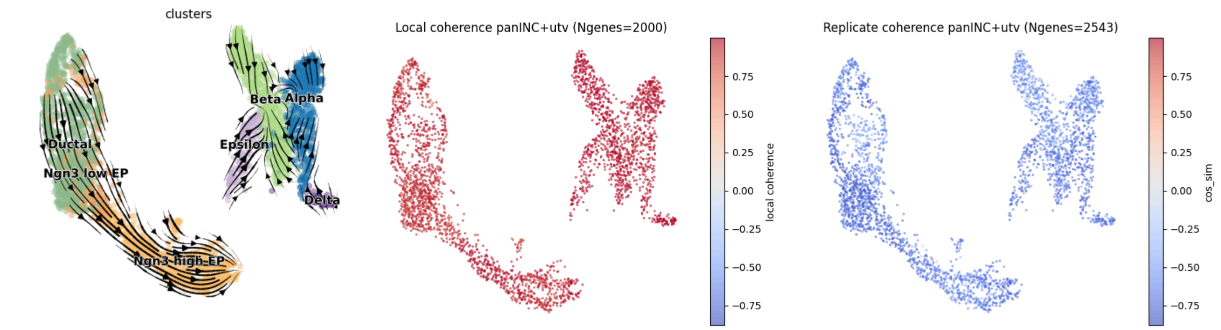


**scTour**

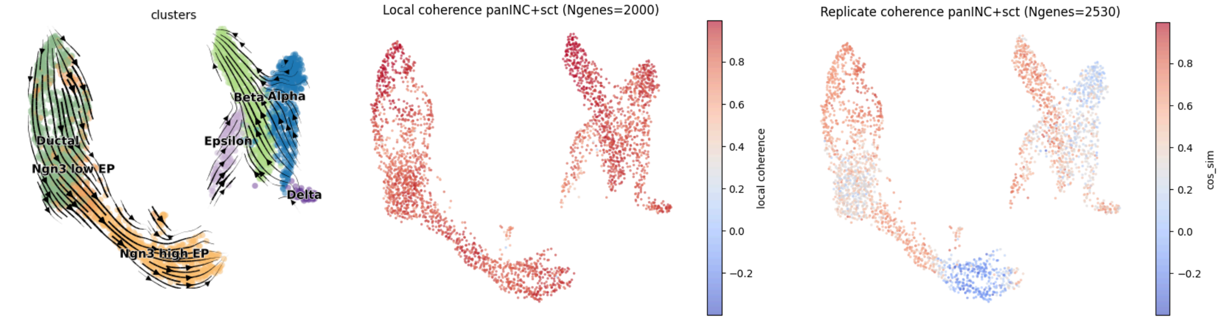


**VeloVI (standard)**

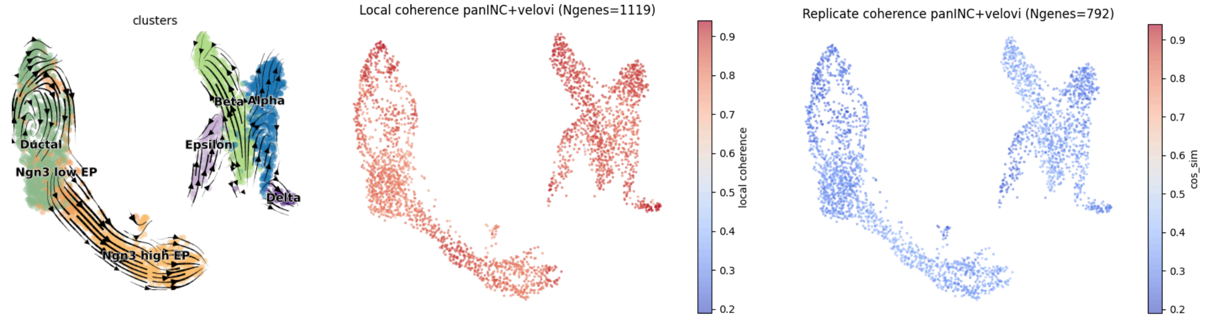


#### VeloVI without preprocessing

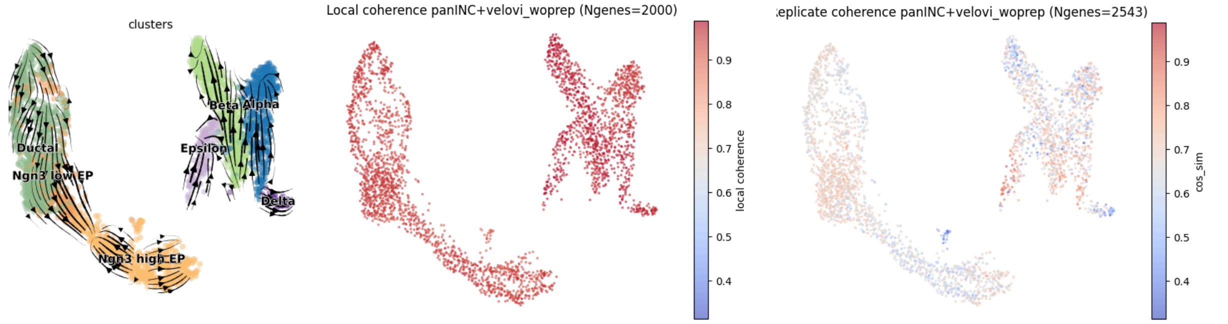


*Figure S14: The same layout as Supplementary Figure S13, except based on analyzing the pancreas dataset without pre-endocrine cells.*

#### <u>UMAPs of brain (using the subset of GPC genes), colored by local and replicate coherence.</u>

#### scVelo

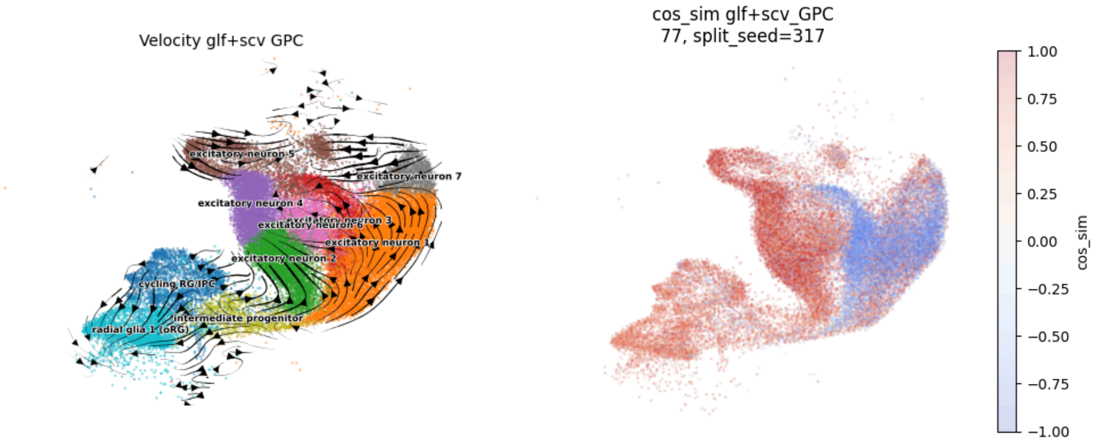


#### UniTVelo

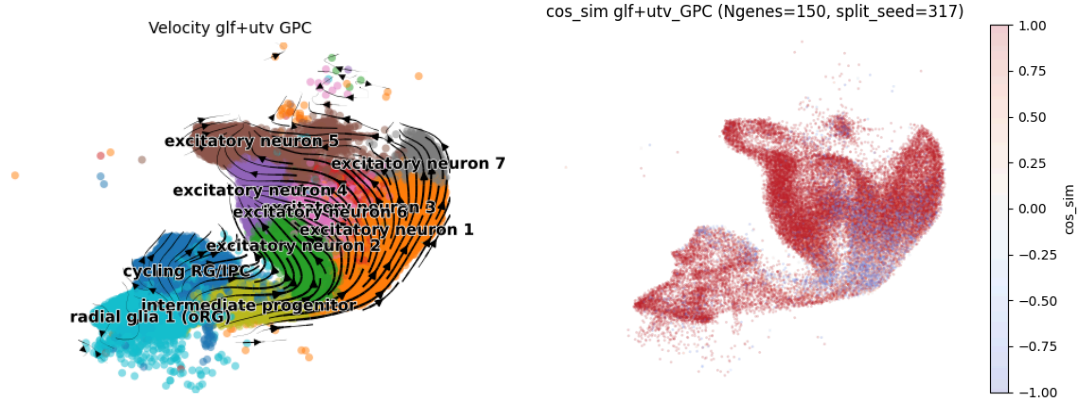


#### scTour

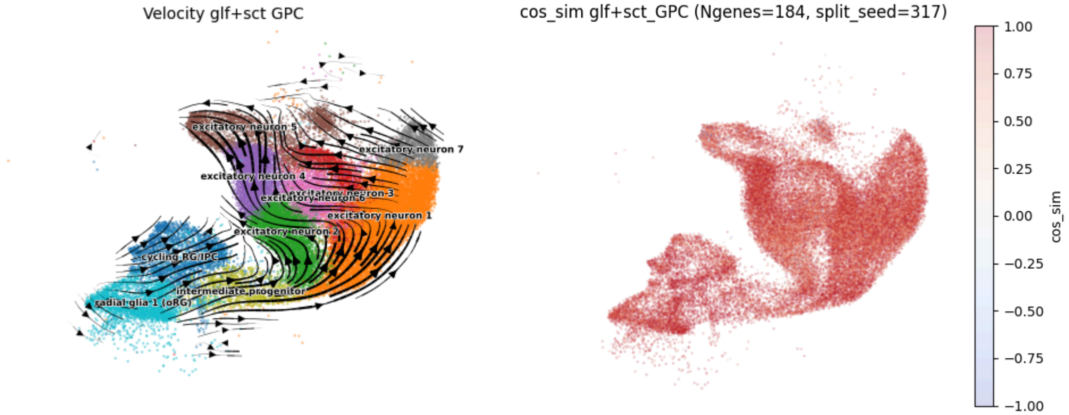


#### VeloVI (standard)

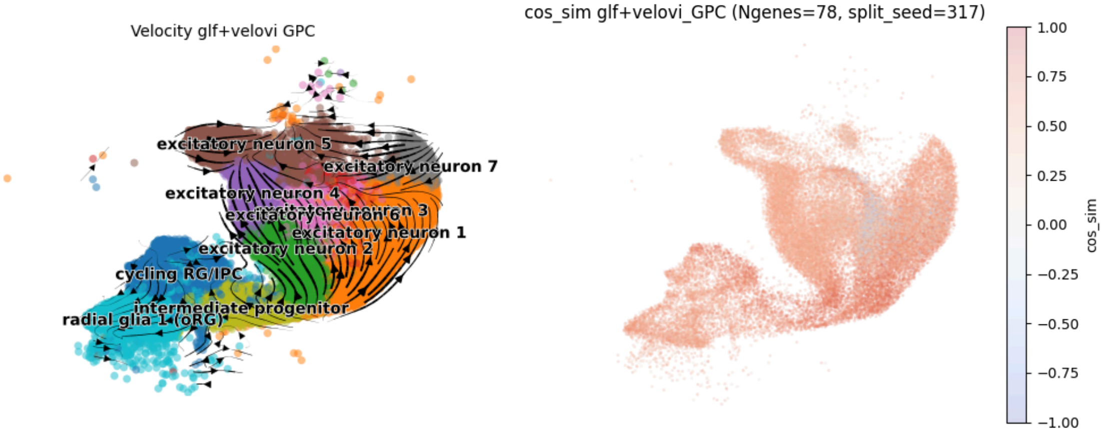


#### VeloVI without preprocessing

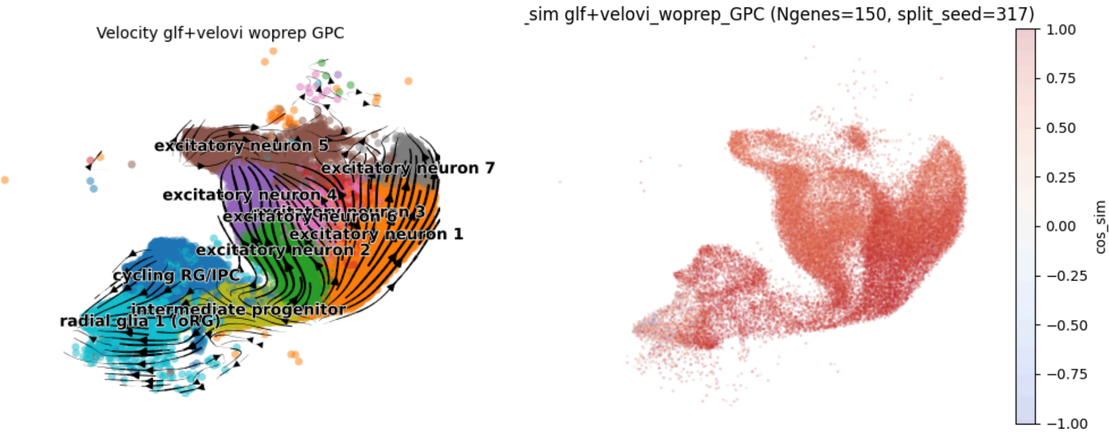


*Figure S15: The replicate coherence (i.e., "velocity cosine similarity", right column) in the brain dataset. The UMAPs with RNA velocity flow are shown in the left column as a reference. The cell type colors are the same as in Figure 4 in the main text, and the same color bar is shown for each of the coherence plots, where red denotes high coherence (i.e., low variability) and blue denotes low coherence (i.e., high variability).*

#### S3.10. ScTour velocity field vs. RNA velocity

As mentioned in Supplemental Section S3.2, scTour natively computes the RNA velocity in a latent space due to its VAE architecture. In this section, we present the following plots, which demonstrate that our procedure for computing the RNA velocity vector (with a dimensionality equal to the number of genes) accurately captures the intended RNA velocity flow. Below is the comparison of the scTour vector field and our recovered velocity estimates, plotted on the original UMAP embedding. The leftmost plot shows each cell type without any velocity estimates. The middle plot shows the vector field produced by scTour. The right plot shows the recovered velocities.

**Erythroid dataset**

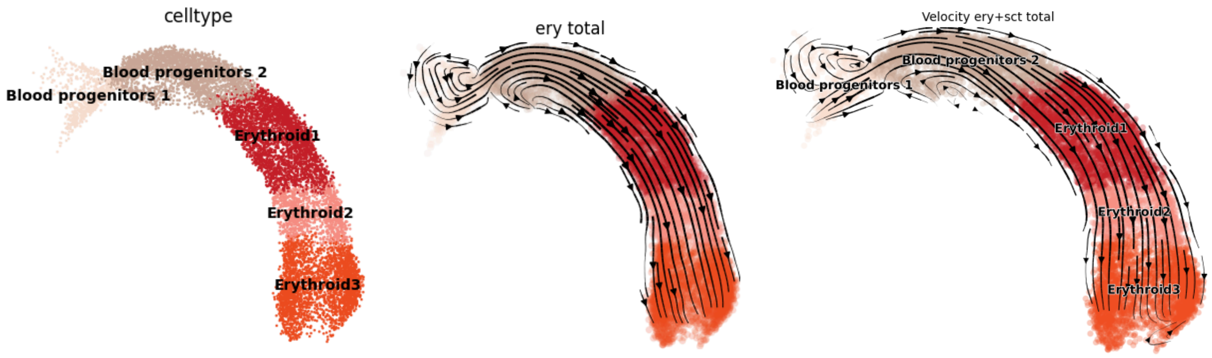


**Pancreas dataset**

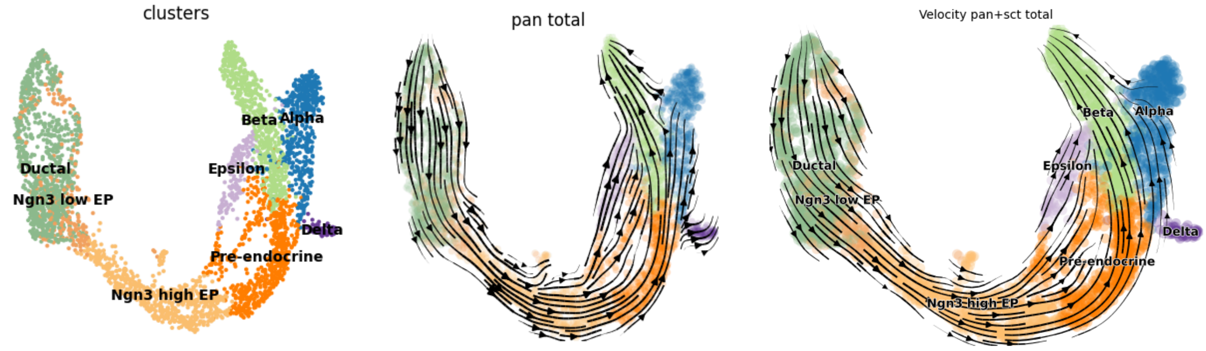


#### Pancreas without pre-endocrine

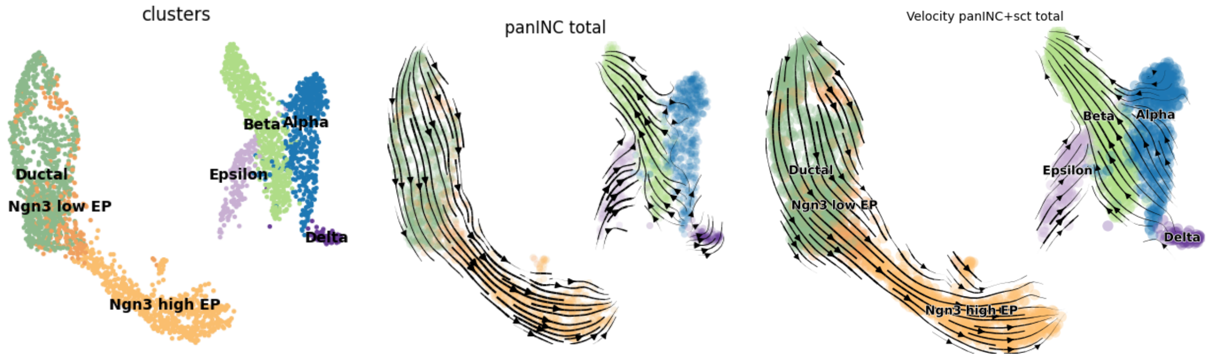


#### Brain dataset

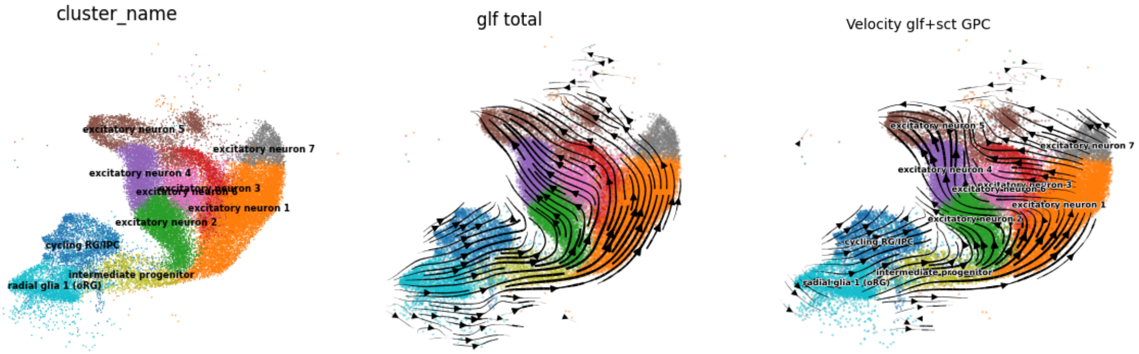


*Figure S16: UMAP comparing the output RNA velocity flow in the latent embedding space estimated by scTour (middle column) and our reconstructed RNA velocity flow across all the genes (right column) for each of the three datasets. The left column displays the UMAP, where the cell type colors match those in Figures 2 through 4 in the main text.*

#### S3.11. Replicate coherence alignment and correlation across random seeds

These plots show the correlations of replicate coherence values across 5 seeds for each RNA velocity method separately. We demonstrate this using the pancreas data with pre-endocrine cells. There are five plots, each showing the correlation between any 2 of the 5 seeds, where we plot each cell by its cell type (also written in the upper-triangular plots).

In each plot, we depict both the “uncentered correlation” (i.e., cosine similarity) between the two sets of replicate coherences, one from each seed, as well as the Pearson correlation. We show the “uncentered correlation” since we want to display whether a positive replicate coherence is indicative of another random seed also yielding a positive replicate coherence. We test for the significance of the uncentered correlation or Pearson correlation via t-tests. We plot the red region to denote when the replicate coherence is negative in either seed – we see that across the 5 RNA velocity methods, very few cells have a negative replicate coherence. This is reassuring, as it demonstrates that the positivity of the replicate coherence is not an artifact of a particular random seed.

#### scVelo

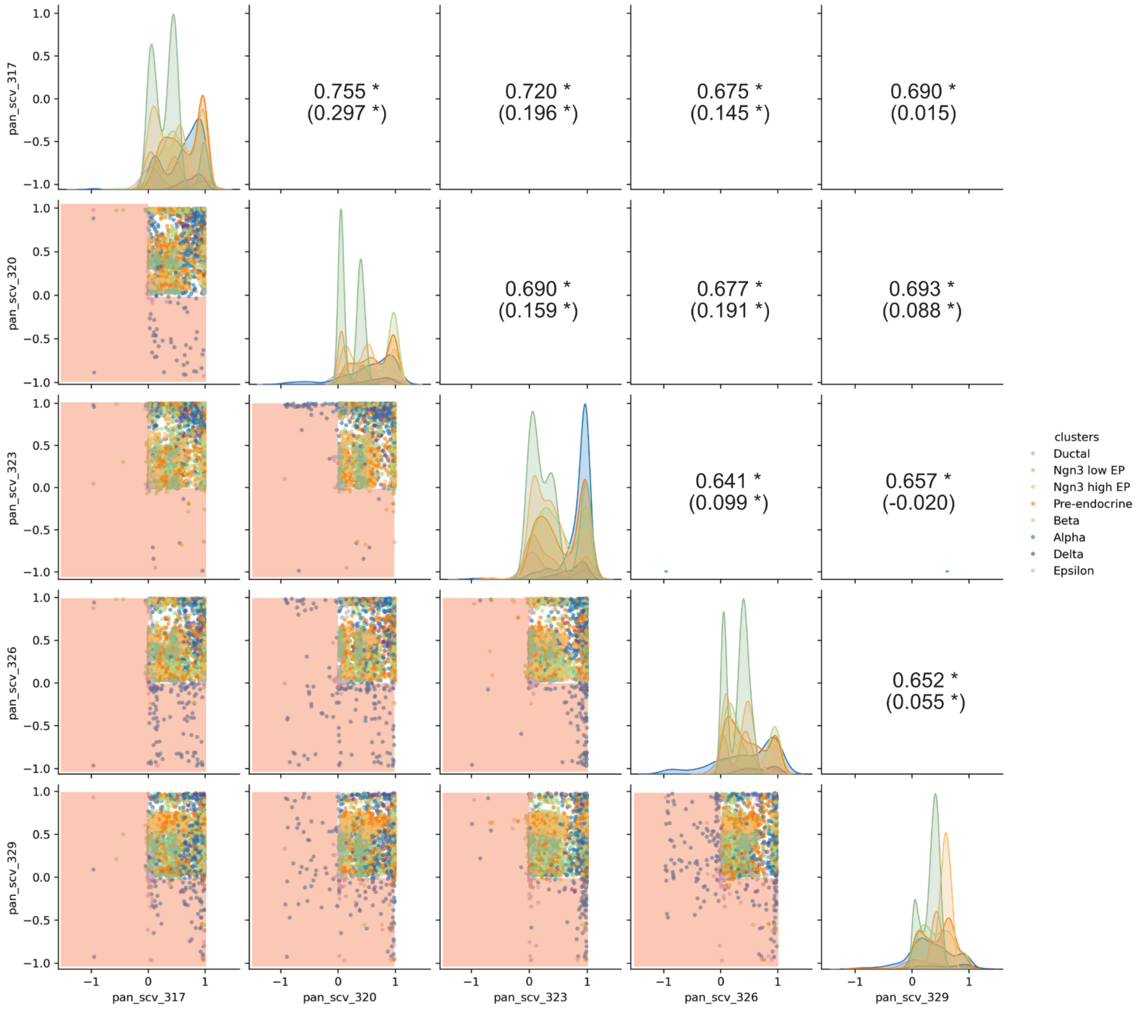


*Figure S17: Pair plot across the five different seeds when applying scVelo on the pancreas dataset. Each point in the lower-triangular plots denotes a cell (colored by cell type), and the x- and y-axes denote the replicate coherence between the two splits for that particular seed. A higher uncentered correlation between the two seeds' replicate coherence denotes more evidence that the replicate coherence based on our framework is not due to randomness inherent to count splitting. We also write the Pearson correlation in parentheses. The red region denotes the region where a cell has a negative replicate coherence (i.e., cosine similarity less than 0) in either seed. We mark statistically significant correlations in the following plots with a * (after multiple testing for correlation), where we test for significance at the level of 0.05. The densities of each seed's replicate coherence are shown along the diagonal, stratified by cell type.*

#### UniTVelo

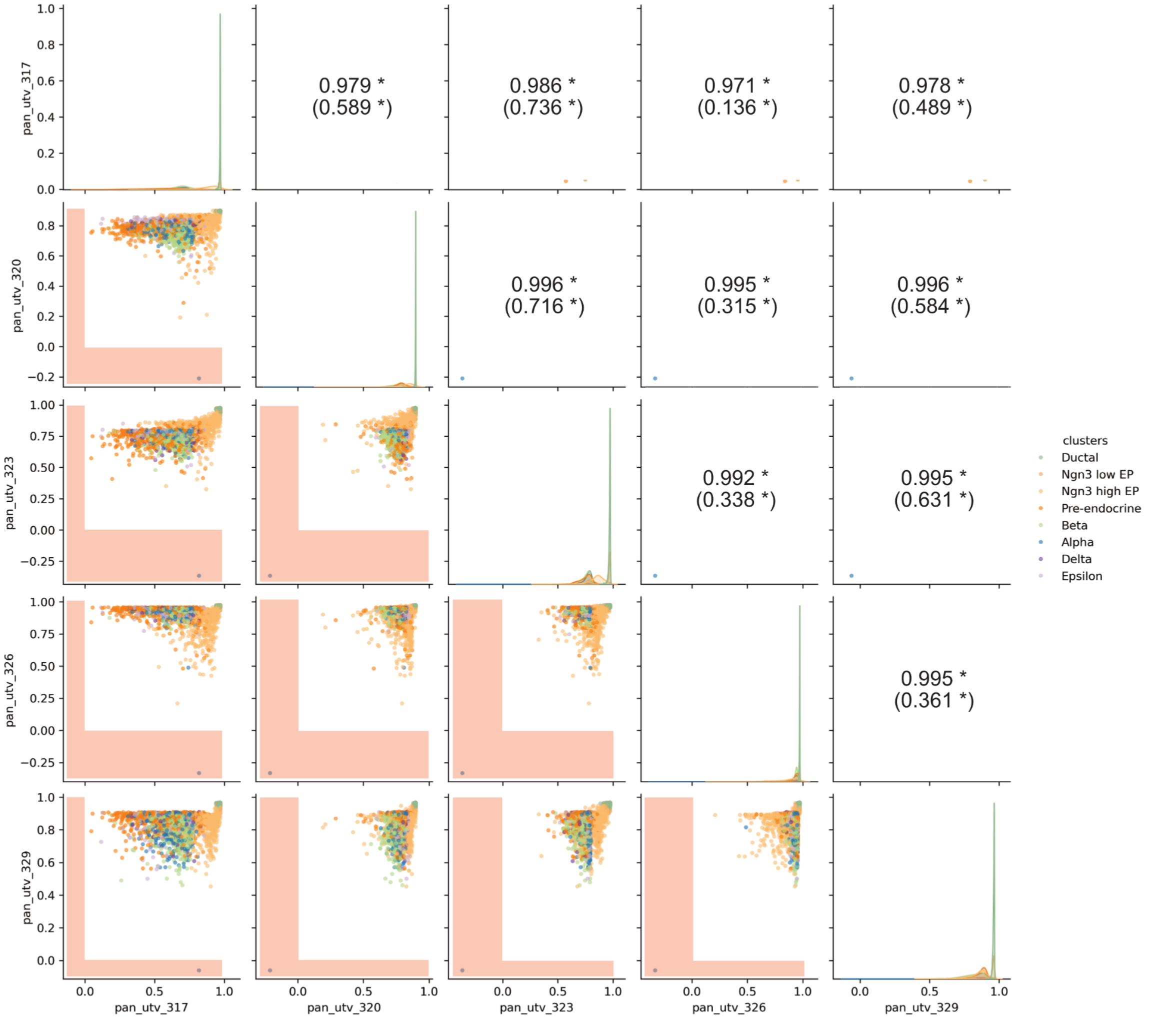


*Figure S18: The same layout as Supplementary Figure S17, except using UniTVelo across five seeds.*

#### scTour

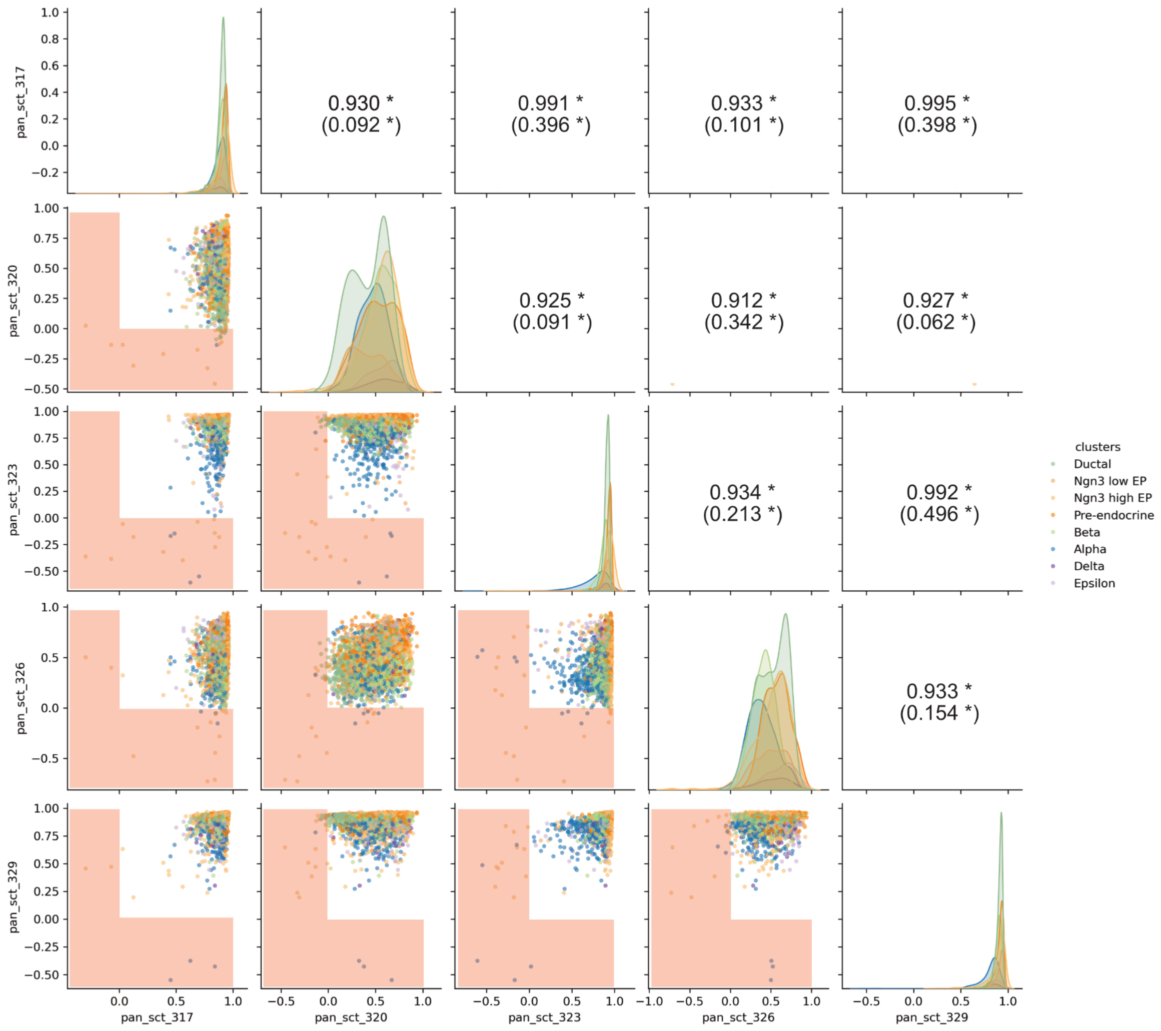


*Figure S19: The same layout as Supplementary Figure S17, except using scTour across five seeds.*

#### Standard VeloVI

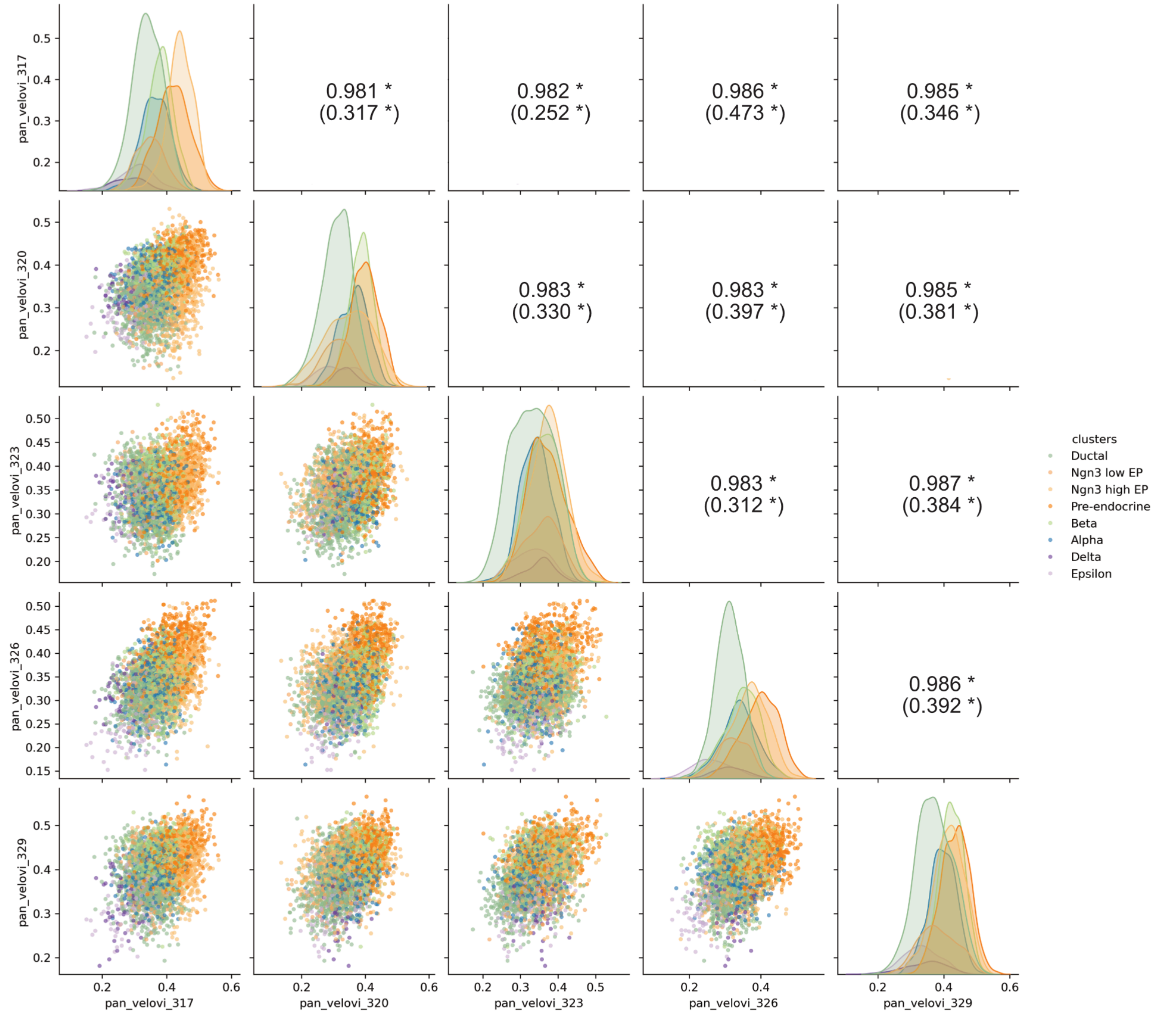


*Figure S20: The same layout as Supplementary Figure S17, except using standard VeloVI across five seeds. No red region is shown since none of the cell's replicate coherence is negative.*

**VeloVI without preprocessing**

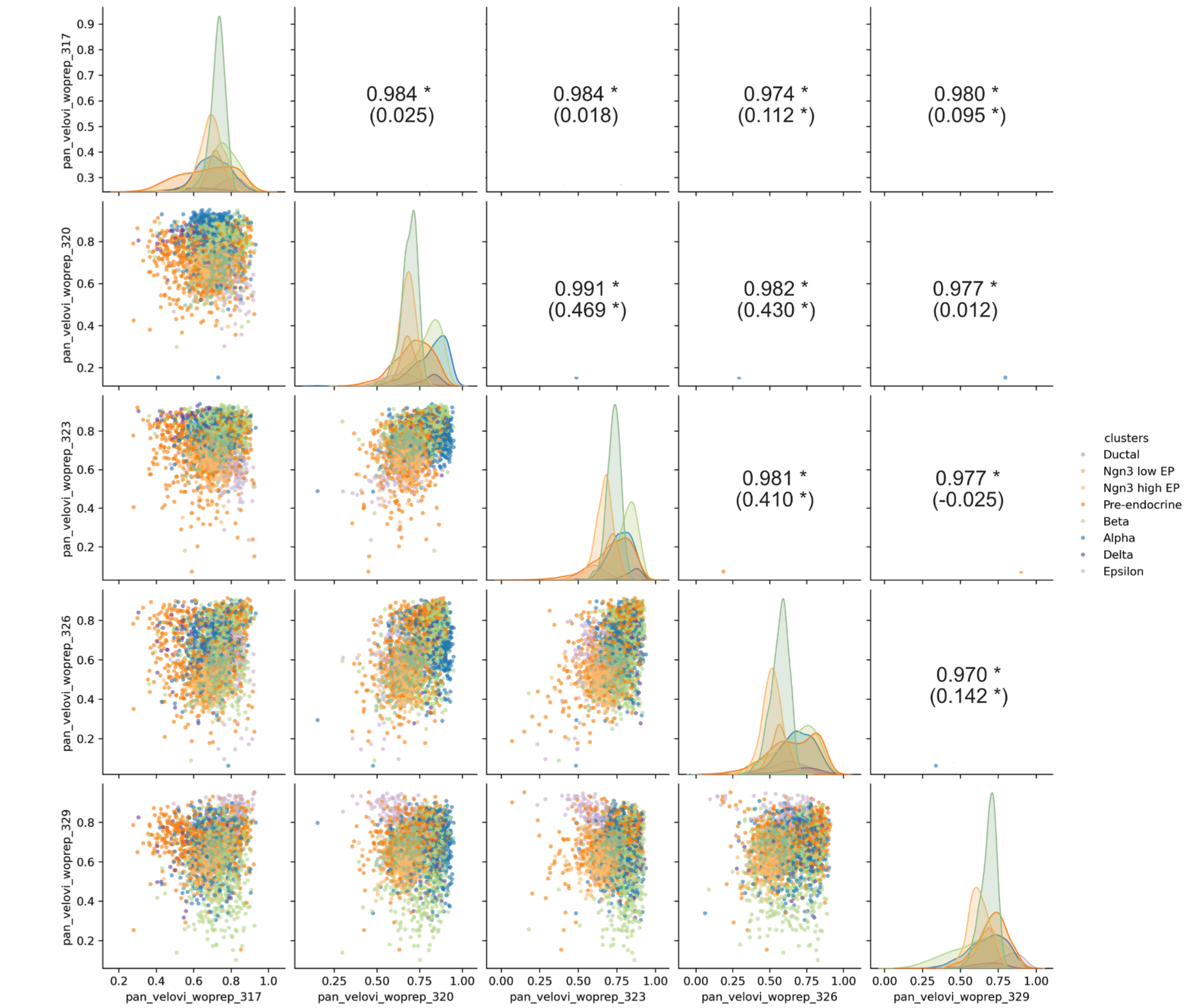


*Figure S21: The same layout as Supplementary Figure S17, except using VeloVI without preprocessing across five seeds. No red region is shown since none of the cell's replicate coherence is negative.*

The following table summarizes the mean and variance of replicate coherence for each method across 5 count splitting runs on the pancreas dataset. The low variance across seeds within each method indicates relatively stable performance, complementing the correlation plots shown in this section.

*Table S5: Mean and variance of replicate coherence across 5 count splitting runs for each method on the pancreas dataset.*

| | scv | utv | sct | velovi (standard) | velovi (w/o prep) |
|---|---:|---:|---:|---:|---:|
| mean(5seeds) | *0.4526* | *0.8469* | *0.7274* | *0.3649* | *0.6883* |
| var(5seeds) | *0.0024* | *0.0034* | *0.0451* | *0.0005* | *0.0026* |

#### S3.12. Computational time of analyses

The table summarizes the computational time, averaged across multiple seeds, for running each method on one split on the two versions of the mouse pancreas data.

*Table S6: Averaged computational time across seeds for running each method on one split on pancreas datasets*

| | **Pancreas data** | **Pancreas data without pre-endocrine** |
|---|---|---|
| *scVelo* | 8min | 7min |
| *UniTVelo* | 238min | 156min |
| *scTour* | 27min | 31min |
| *VeloVI (standard)* | 61min | 56min |
| *VeloVI without preprocessing* | 289min | 255min |

#### S3.13. Definition of informative gene sets

**Erythroid informative genes:** We define the informative genes in the erythroid dataset as the marker genes using Python packages scanpy and scvelo. The procedure involves performing differential expression analysis using the Wilcoxon rank-sum test to compare gene expression between each cell type and all others, to identify genes that are significantly upregulated in specific cell types. The top-ranked genes are selected as marker genes.

**Pancreas informative genes:** The informative genes in the pancreas dataset are retrieved from Table S3 in Bastidas-Ponce et al. (2019), which were the marker genes originally found by the authors. Specifically, those genes are selected based on their expression, by comparing the ductal cluster to Ngn3 low progenitors, Ngn3 low progenitors to the ductal cluster, Ngn3 high precursors to Ngn3 low progenitors, and Fev high cells to Ngn3 high precursors.

**Brain informative genes:** In the brain dataset, the informative genes are selected as those with predictive chromatin (GPC), where their single-cell RNA-seq expression can be predicted by chromatin accessibility measured using single-cell Assay for Transposase-Accessible Chromatin sequencing (scATAC-seq) data after integrating scRNA-seq and scATAC-seq together (Trevino et al., 2021). Specifically, the authors ranked genes based on the correlation between gene activity scores (derived from scATAC-seq) and gene expression (from scRNA-seq). They selected the top 10% of genes with the highest correlations that were also associated with more than 10 putative cis-regulatory elements (CREs). This yielded a set of 185 GPC genes, which were notably enriched for transcription factors and DNA-binding regulators, suggesting they play key roles in establishing cell identity during cortical development. These GPCs were further validated using multiomic data -- simultaneous RNA and chromatin profiling from the same cells -- which confirmed that the majority of CRE-gene links were preserved and that GPC expression-accessibility correlations held at the single-cell level.

#### S3.14. Signal-to-random coherence score for erythroid and pancreas dataset

This section presents the summary of signal-to-random coherence scores obtained from the five RNA velocity methods for the erythroid and pancreas datasets. The scores are displayed as boxplots for method comparison, and the exact values are provided in tables following each plot.

**Erythroid dataset**

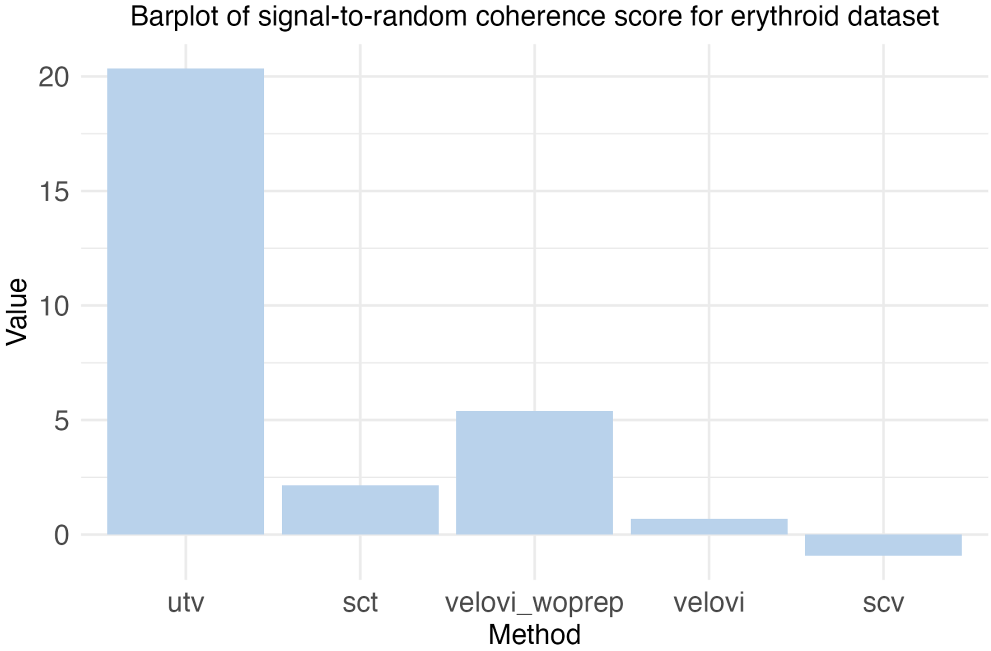


*Figure S22: Barplot of the signal-to-random coherence score across the five RNA velocity methods for the erythroid dataset.*

*Table S7: Signal-to-random coherence scores across the five RNA velocity methods for the erythroid dataset.*

| utv | sct | velovi_woprep | velovi | scv |
|---|---|---|---|---|
| 20.3409 | 2.1411 | 5.3902 | 0.6875 | -0.9136 |

**Pancreas dataset**

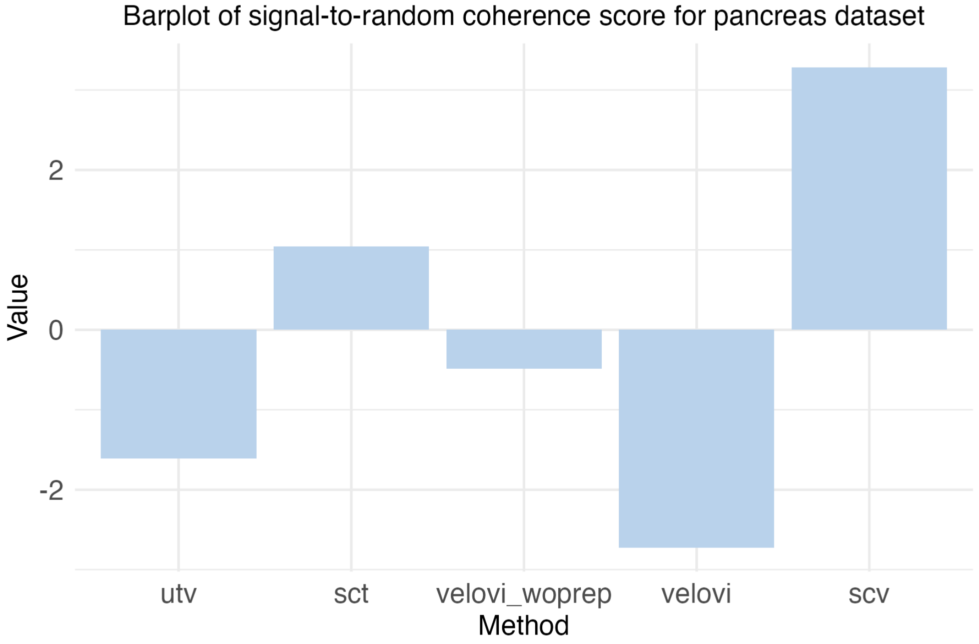


*Figure S23: Barplot of the signal-to-random coherence score across the five RNA velocity methods for the pancreas dataset.*

*Table S8: Signal-to-random coherence scores across the five RNA velocity methods for the pancreas dataset.*

| utv | sct | velovi_woprep | velovi | scv |
|---|---|---|---|---|
| -1.6089 | 1.0418 | -0.4873 | -2.7230 | 3.2812 |

#### S3.15. Identifying genes aligned with direction-aware Laplacian transition matrix

We apply CellRank's velocity kernel to compute a transition matrix based on an RNA velocity method's results (i.e., the velocity vector and gene expression of every cell). This transforms every cell into a probability-weighted mixture of its putative future neighbours. We view it as a directed graph whose edges encode the flow implied by RNA velocity. A gene whose expression pattern faithfully follows this flow should vary smoothly along these directed edges – i.e., large expression differences should be rare on edges with high transition probability. Conversely, genes whose expression is orthogonal to the velocity field will fluctuate "against the current" and therefore appear rough on the same graph.

The Chung-directed Laplacian furnishes an energy functional that quantifies precisely this notion of smoothness. It's Rayleigh quotient, here called the *Laplacian score*, assigns one scalar to every gene:

- a small score means the expression is coherent with the velocity-defined transitions, and
- a large score means expression fluctuates across high-probability transitions.

Ranking genes by this score and feeding the ranking into GSEA enables us to discover biological processes whose transcriptomic programs are most tightly coupled to the inferred developmental dynamics.

We now describe the details:

**Step 1: Transition matrix.**

Let $A \in \mathbb{R}_{\geq 0}^{n \times n}$ be the sparse velocity-graph adjacency returned by Cellrank. A row normalisation yields the Markov matrix $P \;=\; D_{\text{out}}^{-1} A$ where $D_{\text{out}} \;=\; \text{diag}(A\mathbf{1})$, so that $(P_{ij} = \Pr(i \to j)$ and $\sum_j P_{ij} = 1$.

**Step 2: Chung-directed Laplacian**

We next compute the stationary distribution $\pi^\top P = \pi^\top$ via power iteration. Define $S = D_{\sqrt{\pi}}\, P\, D_{1/\sqrt{\pi}}$ where $D_{\sqrt{\pi}} = \text{diag}(\sqrt{\pi_1}, \ldots, \sqrt{\pi_n})$. Then, the Chung-directed Laplacian is defined as $L \;=\; I - H$ for $H = (S + S^\top)/2$. Here, $L$ is positive-semidefinite and coincides with the usual random-walk Laplacian when the graph is undirected (Chung, 2005).

**Step 3: Gene-level Laplacian score**

For every gene $g$, let $x^{(g)} = (x_1^{(g)}, \ldots, x_n^{(g)})$ be its (spliced) expression. The $\pi$-weighted mean and variance are $\mu^{(g)} = \sum_{i=1}^{n} \pi_i\, x_i^{(g)}$ and $\text{Var}_\pi(\mathbf{x}^{(g)}) = \sum_{i=1}^{n} \pi_i (x_i^{(g)} - \mu^{(g)})^2$. We define the Laplacian score as the typical Laplacian score, but normalized by the $\pi$-weighted variance:

$$\text{LS}(g) = \frac{(x^{(g)\top} L x^{(g)}}{\text{Var}_\pi(\boldsymbol{x}^{(g)})} = \frac{\sum_{i,j} \pi_i\, P_{ij} (x_i^{(g)} - x_j^{(g)})^2}{2\text{Var}_\pi(x^{(g)})}$$

Small values indicate that the gene varies smoothly along high-probability velocity transitions, whereas large values denote roughness. Genes with $\text{Var}_\pi(\mathbf{x}^{(g)}) \approx 0$ are discarded.

**Step 4: Gene-set enrichment**

Because stronger alignment corresponds to smaller $\text{LS}(g)$, we rank genes by $t_g = 1/\text{LS}(g)$ in decreasing order and perform preranked GSEA with clusterProfiler::gseGO (ontology = BP, 10 ≤ size ≤ 500, FDR threshold 0.05). Significant pathways pinpoint biological processes whose transcriptomic programmes propagate coherently with the RNA-velocity field.

### S3.16. Biological pathways aligned with RNA velocity field chosen by the signal-to-random coherence score

In this subsection and the main text, we only focus on pathways with: 1) a p-value less than 0.05, and 2) a core enrichment with more than 5 genes. This is to filter out pathways that are not statistically aligned with the direction-aware Laplacian transition matrix, or are driven by too few genes for biological interpretation. The following results are based on the selected RNA velocity method with the highest signal-to-random score for their respective datasets (see Supplementary Section S3.15).

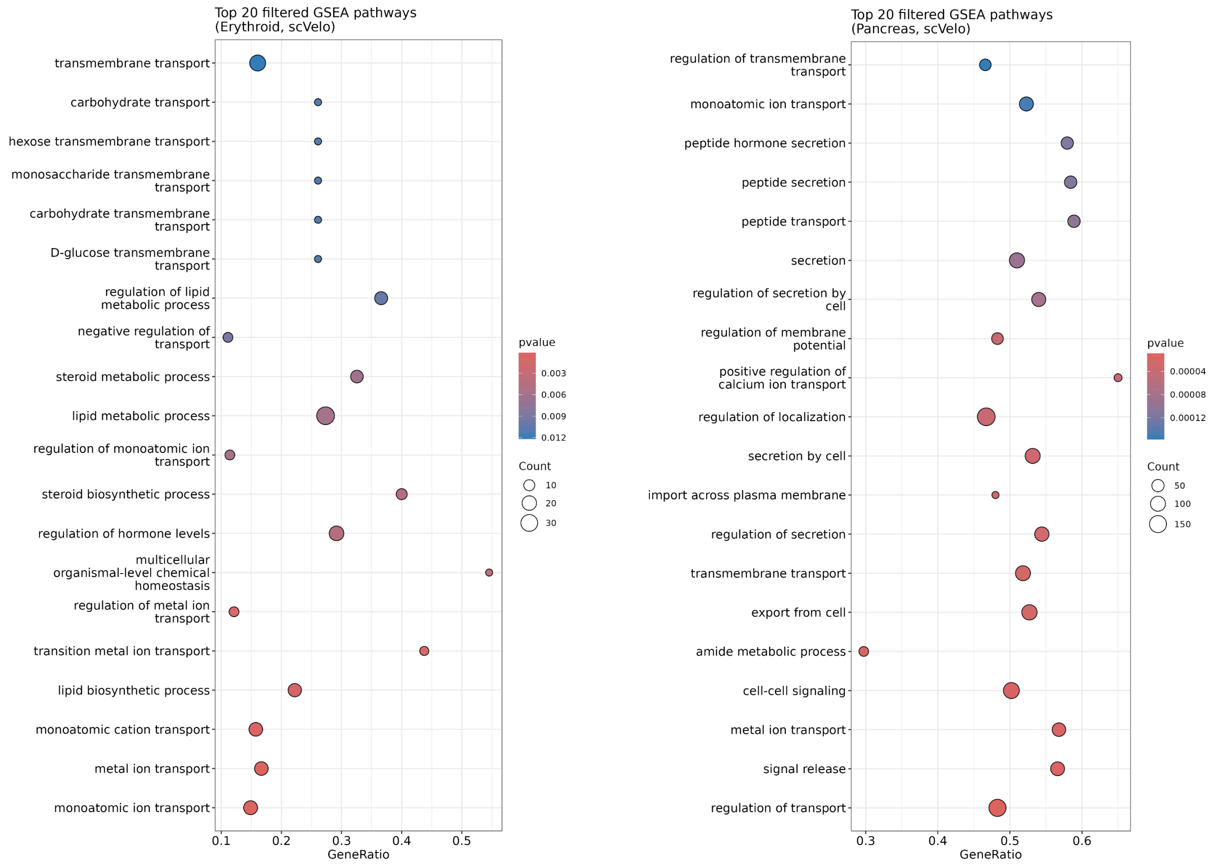


*Figure S24: Dotplot of the enriched pathways that align with the RNA velocity vector field. Both the enriched pathways for the erythroid data (via UniTVelo, left) and pancreas dataset (via scVelo, right) are shown. We choose these specific methods because they have the highest signal-to-random score in their respective datasets (see Supplementary Figure S22 and Figure S23).*

We also show the enriched pathways for the human developing brain dataset, where we show the pathways based on veloVI without preprocessing fit based on the alignment across all the cells in the entire dataset (not just the Excitatory Neuron 4).

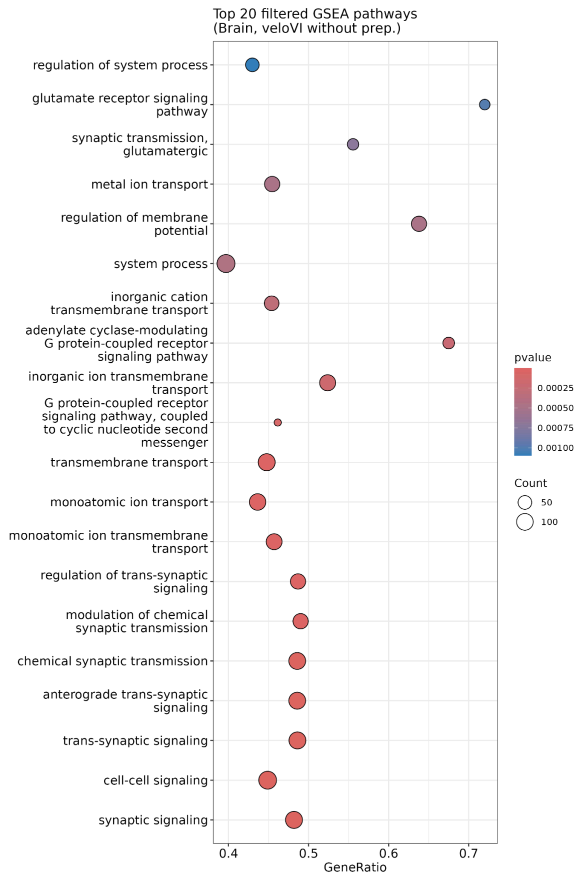


*Figure S25: Dotplot of the enriched pathways that align with the RNA velocity vector field for the brain dataset using the veloVI without preparation method, which has the highest signal-to-random score on this dataset. These results are based on RNA velocity fitting for all cells, not just the Excitatory Neuron 4 cells.*

#### S3.17. Shuffled cosine similarity on pancreas

It is possible that in some cases, the high cosine similarity in replicate coherence (either in terms of a summary statistic or the overall distribution) is likely to result from the similarity of the velocity vectors across all cells, rather than the “true” similarity between cells in one split and their ‘twins’ in another.

This can be viewed through a bias-variance lens. For example, a degenerate RNA velocity method that ignores the data and returns the same velocity vector for every cell (e.g., an all-ones vector) would achieve replicate coherence 1 under count splitting (perfect stability), yet be uninformative and generally inaccurate. Thus, replicate coherence primarily reflects the *variance* of the estimator across splits. To probe the complementary failure mode (stable but overly similar velocities across cells), we could compare a cell’s twin-pair similarity to similarity with a randomly paired cell via the shuffled-pair procedure below.

To account for the potential effect of velocity vector similarity across cells, we apply the following procedure: as our procedure already prescribes, we first count split the spliced and unspliced count matrices and apply RNA velocity on each split. We then compute each cell's replicate coherence ("paired"). Then, we suggest the new steps afterwards: we fix the velocity vectors in one split and shuffle the order of cells in another split. Then, we recompute the cosine similarity of velocity vectors between the shuffled cell pairs, which can quantify how similar a cell's velocity vector is compared to a random cell in another split ("shuffled"). Importantly, in these new steps, no additional RNA velocity estimation is performed.

If the replicate coherence computed on the paired cells is higher than that of the shuffled cells and the difference is small, we may consider the performance unstable, even if the value is close to 1. Based on this idea, a larger difference in cosine similarity between paired and shuffled cell pairs indicates a more non-random behavior.

Below is a summary of the paired and shuffled mean and median cosine similarity values for pancreas cells between the two splits for each method. The last two rows ("diff mean" and "diff median") are the differences in the paired and shuffled values. Here, a larger difference means that there is a larger distinction between the cell's original estimated RNA velocity vector and a randomly assigned RNA velocity vector.

*Table S9: Replicate coherence (i.e., "paired") summarized as mean and median across all cells, as well as the replicated coherence after shuffling the RNA velocity vectors (i.e., "shuffled"), and the difference between the two, for each of the five RNA velocity methods*

| | scVelo | UniTVelo | scTour | VeloVI (std) | VeloVI (w/o prep) |
|---|---|---|---|---|---|
| paired mean | 0.492 | 0.774 | 0.8446 | 0.378 | 0.7122 |
| paired median | 0.4658 | 0.7727 | 0.871 | 0.3786 | 0.7225 |
| shuffled mean | 0.41013 | 0.2575 | 0.11892 | 0.199 | 0.4052 |
| shuffled median | 0.2769 | 0.4055 | -0.02414 | 0.1688 | 0.4139 |
| diff mean | 0.08187 | 0.5165 | 0.72568 | 0.179 | 0.307 |
| diff median | 0.1889 | 0.3672 | 0.89514 | 0.2098 | 0.3086 |

However, we note that this shuffled comparison does not account for method-specific properties, such as differences in the smoothness or regularization of the estimated velocity vector field. Without adjusting for these aspects, the raw difference between paired and shuffled cosine similarities may offer an overly simplistic assessment of stability.

#### S3.18. Simulations to demonstrate premise of replicate coherence

In this section, we demonstrate the premise of replicate coherence via simulation where we show that if there's a higher replicate coherence (i.e., higher mean cosine similarity between Split 1 and Split 2), this data-driven result is suggestive of a higher accuracy of the velocity field (i.e., higher mean cosine similarity between an RNA velocity analysis on the entire dataset and the "true" velocity).

To generate the data, we use the `scvelo.datasets.simulation` function in the scvelo package, where we set `n_obs = 1000` (i.e., the number of cells), `n_vars = 50` (i.e., the number of genes), `t_max = 25`, `alpha = 5`, `beta = .3`, `gamma = .5`, and `noise_level = 0.1`. We use this simulation function since it generates each cell's transcriptomic profile via the ODE RNA splicing framework (i.e., a correctly specified model), and we use a relatively small noise level to make our simulation transparent. When we applied this simulation across multiple trials and with various simulation settings, the overall conclusions remain the same.

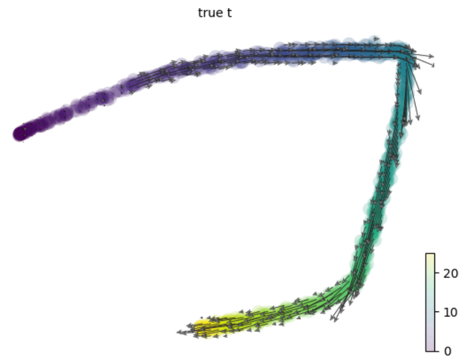


*Figure S26: The UMAP of the simulated data (cells colored by pseudotime) with the true velocity (defined as β*[Unspliced counts] - γ*[Spliced counts]).*

In the simulation study we show below, we focus on both versions of scVelo (stochastic and dynamical mode), as the simulated data from the scVelo package is most compatible with scVelo estimation procedure. Since different RNA velocity methods define RNA velocity slightly differently, using other RNA velocity methods (e.g., VeloVI or UniTVelo) yields an RNA velocity vector that is not easily comparable to the "true" RNA velocity vector simulated via the scVelo package due to the different "units" each method uses.

**scVelo: Stochastic mode**

We first apply scVelo using 'stochastic mode' (i.e., analogous to Velocyto) on the data using all the counts.

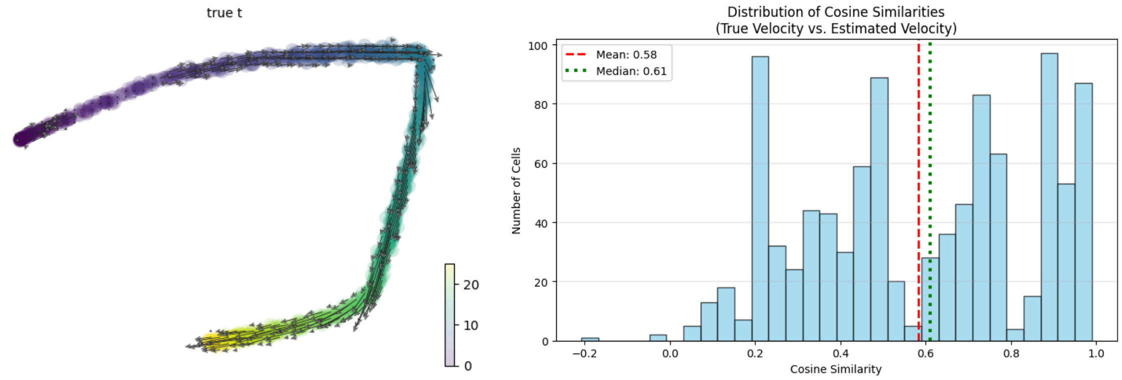


*Figure S27: (Left) UMAP showing the estimated RNA velocity vectors using scVelo's stochastic mode, where the UMAP coordinates are the same as those in Figure S26. (Right) The cosine similarity of each cell, comparing its estimated and true velocity vectors. The mean and median values are marked.*

As prescribed in our procedure, we then estimate the overdispersion of each gene and apply count splitting. When applying scVelo in stochastic mode to each split, we get the following visualizations. We note that the UMAPs for Split 1 and Split 2 (Figure S28) individually are a lot more "dispersed" than the UMAP of the total counts (Figure S27, left). This demonstrates that each split has "less information" than the total counts. This is analogous to cross-validation for hyperparameter tuning in machine learning (e.g., tuning Lasso or XGBoost), where splitting a dataset into different folds decreases the sample size. The assessment based on the "smaller data" is then used to inform how to analyze the "full dataset." Likewise, we see that in our count splitting framework, there is still ample information to measure the coherence of RNA velocity vectors between the two splits.

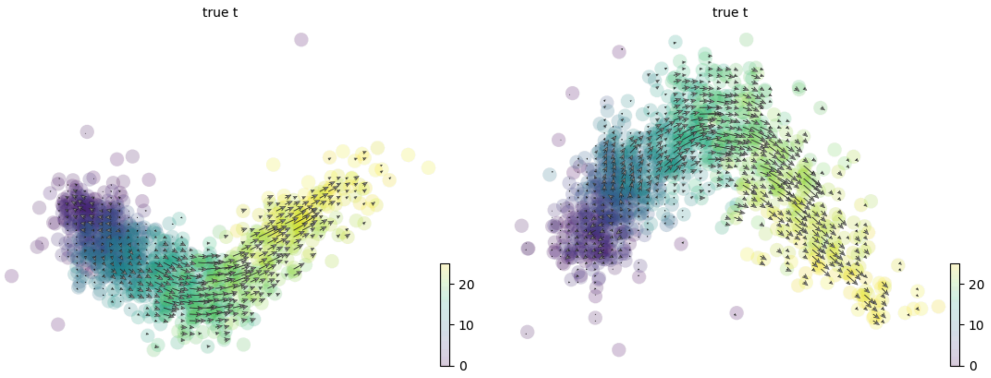


*Figure S28: (Left) UMAP of Split 1 showing the estimated RNA velocity vectors using scVelo's stochastic mode. (Right) Same, except for Split 2.*

Finally, comparing the cosine similarity between each cell's velocity vector between Split 1 and Split 2, we observe a replicate coherence of about 0.4.

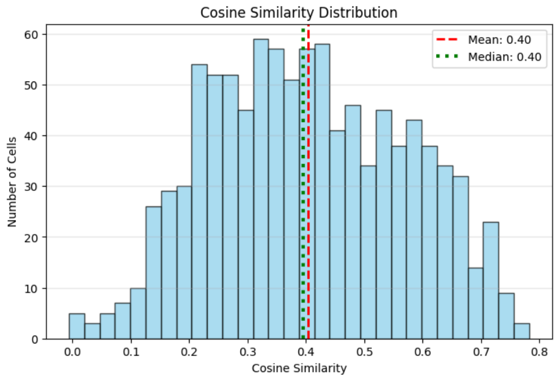


*Figure S29: The cosine similarity of each cell, comparing its velocity vectors (scVelo stochastic mode) on Split 1 and Split 2. The mean and median values are marked.*

<u>An important note</u> is that we are <u>not</u> advocating for using the RNA velocity vectors estimated on each split for biological investigation. The RNA velocity vectors estimated from each split individually are typically less accurate compared to the RNA velocity vectors estimated from the total count data. To reiterate: the premise of our work is that the replicate coherence (i.e., stability) of the RNA velocity vectors between the two splits is an important quality needed for accurate RNA velocity vectors estimated on the total data.

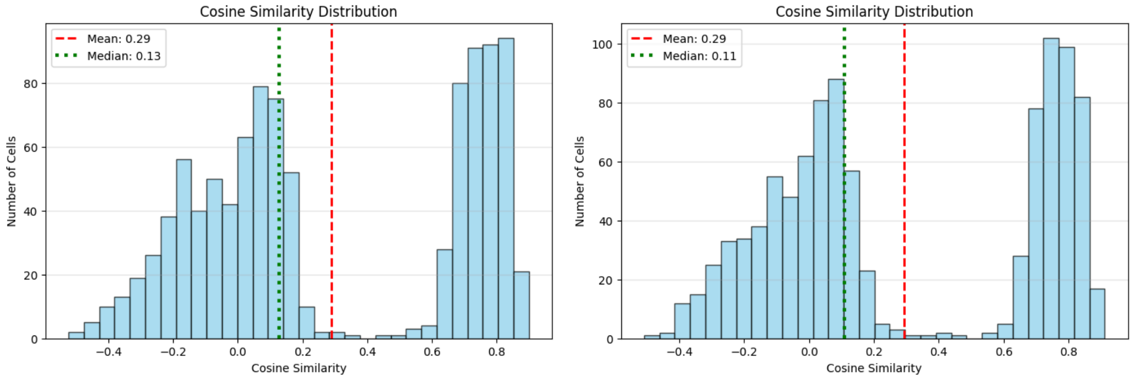


*Figure S30: The cosine similarity of each cell, comparing its velocity vectors (scVelo stochastic mode) estimated on each split (left: Split 1, right: Split 2) to the true velocity vectors. The mean and median values are marked. Comparing these plots to Figure S27 (right) demonstrates that the RNA velocity vector of each individual split is not accurate, even though the RNA velocity vector of the total data is accurate.*

**scVelo: Dynamical mode**

In order to facilitate a comparison, we also analyze the data using scVelo in ‘dynamical mode’ on the data using all the counts.

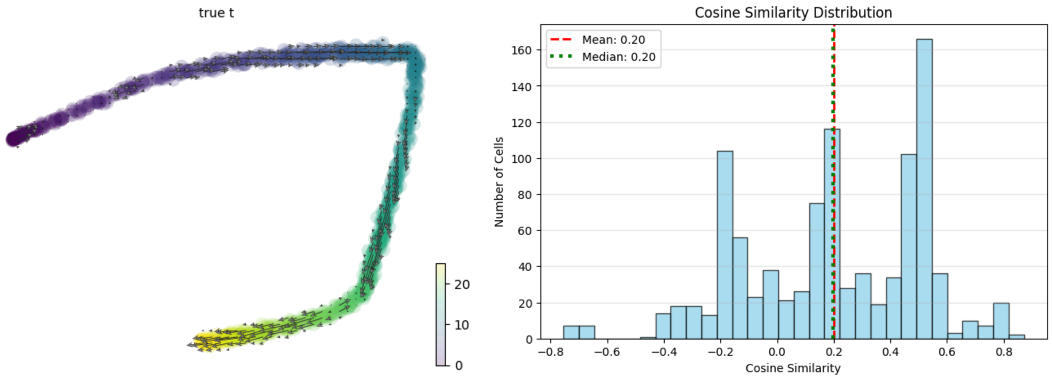


*Figure S31: (Left) UMAP showing the estimated RNA velocity vectors using scVelo's dynamical mode, where the UMAP coordinates are the same as those in Figure S26. (Right) The cosine similarity of each cell, comparing its estimated and true velocity vectors. These plots are analogous to Figure S27.*

Similar to our aforementioned analysis, we use the same count-splitted data and apply scVelo (dynamical mode) on Split 1 and Split 2.

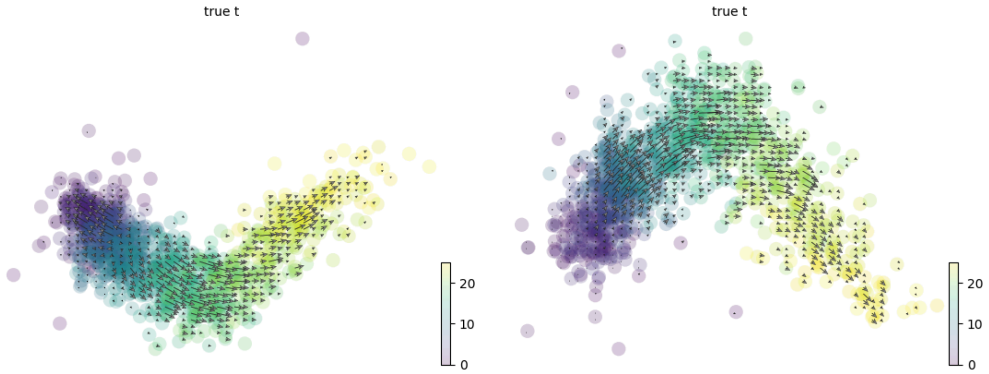


*Figure S32: (Left) UMAP of Split 1 showing the estimated RNA velocity vectors using scVelo's dynamical mode. (Right) Same, except for Split 2. These plots are analogous to Figure S28.*

Finally, we compute the replicate coherence for scVelo's dynamical mode. We see the replicate coherence is 0.19, which is lower than that of scVelo in the stochastic model. This demonstrates our premise: The lower replicate coherence (comparing Figure S33 to Figure S29) is suggestive of the accuracy of the RNA velocity fit (comparing Figure S31 right to Figure S27 right).

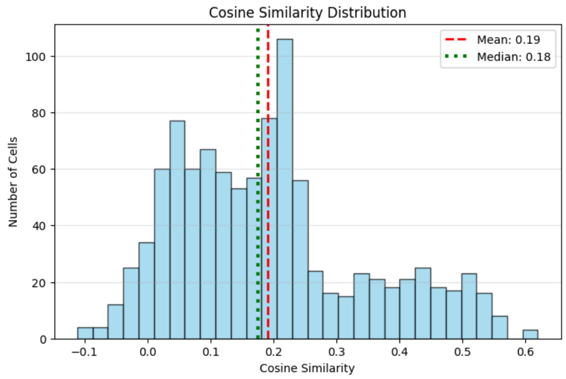


*Figure S33: The cosine similarity of each cell, comparing its velocity vectors (scVelo dynamical mode) on Split 1 and Split 2. This plot is analogous to Figure S29.*

**Simulation over 50 trials:**
Equipped with these results, we then iterated this over 50 trials. Each trial uses the same data generation strategy via scVelo's simulation function, but yields different spliced and unspliced count matrices. In over 50 trials, the replicate coherence for the stochastic mode was higher than the dynamical mode.

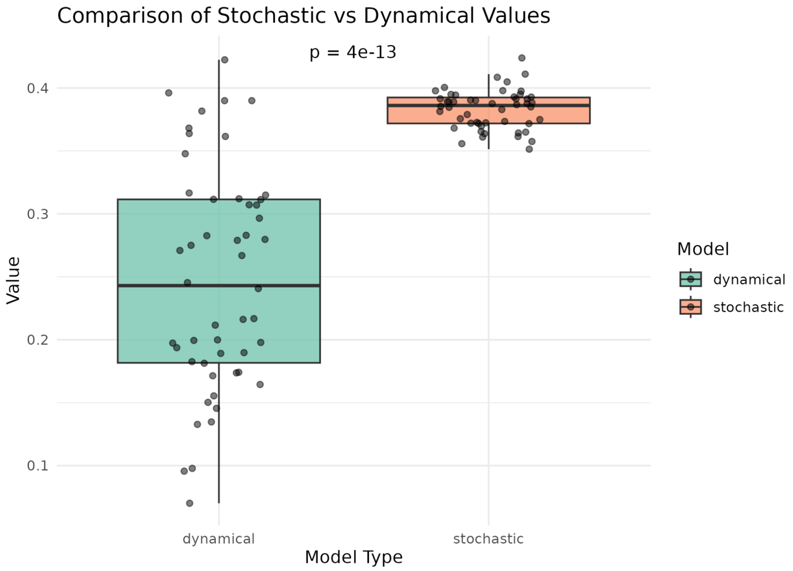


*Figure S34: Replicate coherence scores across 50 trials using scVelo in stochastic or dynamical mode. The p-value shown is using a two-sided Wilcoxon rank-sum test, demonstrating that the stochastic mode's replicate coherence is statistically significantly higher than that for the dynamical mode.*

Comprehensive single cell mRNA profiling reveals a detailed roadmap for pancreatic endocrinogenesis. *Development*, *146*(12), dev173849. https://doi.org/10.1242/dev.173849

- Bergen, V., Lange, M., Peidli, S. *et al.* Generalizing RNA velocity to transient cell states through dynamical modeling. *Nature Biotechnology* 38, 1408–1414 (2020). https://doi.org/10.1038/s41587-020-0591-3
- Chung, F. (2005). Laplacians and the Cheeger inequality for directed graphs. *Annals of Combinatorics*, *9*, 1-19. https://doi.org/10.1007/s00026-005-0237-z
- Gao, M., Qiao, C. & Huang, Y. UniTVelo: temporally unified RNA velocity reinforces single-cell trajectory inference. *Nature Communications* 13, 6586 (2022). https://doi.org/10.1038/s41467-022-34188-7
- Gayoso, A., Weiler, P., Lotfollahi, M. *et al.* Deep generative modeling of transcriptional dynamics for RNA velocity analysis in single cells. *Nature Methods* 21, 50–59 (2024). https://doi.org/10.1038/s41592-023-01994-w
- La Manno, G., Soldatov, R., Zeisel, A., *et al.* RNA velocity of single cells. *Nature*, 560(7719):494–498, (2018).
- Li, Q. scTour: a deep learning architecture for robust inference and accurate prediction of cellular dynamics. *Genome Biology* 24, 149 (2023). https://doi.org/10.1186/s13059-023-02988-9
- Pijuan-Sala, B., Griffiths, J.A., Guibentif, C. *et al.* A single-cell molecular map of mouse gastrulation and early organogenesis. *Nature* 566, 490–495 (2019). https://doi.org/10.1038/s41586-019-0933-9
- Polański, K., Young, M. D., Miao, Z., Meyer, K. B., Teichmann, S. A., & Park, J. E. (2020). BBKNN: Fast batch alignment of single cell transcriptomes. *Bioinformatics*, *36*(3), 964-965.
- A. E. Trevino, F. Müller, J. Andersen, L. Sundaram, A. Kathiria, A. Shcherbina, K. Farh, H. Y. Chang, A. M. Pas, ca, A. Kundaje, et al. Chromatin and gene-regulatory dynamics of the developing human cerebral cortex at single-cell resolution. *Cell*, 184(19):5053–5069, 2021.
